# Combinatorial neural inhibition for stimulus selection across space

**DOI:** 10.1101/243279

**Authors:** Nagaraj R. Mahajan, Shreesh P. Mysore

## Abstract

The ability to select the most salient stimulus among competing ones is essential for animal behavior and operates regardless of the spatial locations that stimuli occupy. Here, we reveal that the brain employs a combinatorially optimized strategy to solve such location-invariant stimulus selection. With experiments in a key inhibitory nucleus in the vertebrate midbrain selection network, called isthmi pars magnocellularis (Imc) in owls, we discovered that the central element is a ‘multilobe’ neuron, which encodes visual locations with multiple firing fields. This multilobed coding of space is necessitated by scarcity of Imc neurons. Although distributed seemingly randomly in space, the locations of these lobes are optimized across the high firing Imc neurons, allowing them to cooperatively suppress stimuli throughout 2D visual space while minimizing metabolic and circuit wiring costs. Our work suggests that combinatorial coding of space by sparse inhibitory neurons may be a general functional module for spatial selection.

---

Animals routinely encounter multiple competing pieces of information in their sensory environments. Typically, they handle this informational complexity by selecting the most salient or behaviorally relevant piece of information, i.e., highest ‘priority’ information, to guide their actions ^1–3^. However, how neural circuits orchestrate the computations that are essential for such stimulus selection is not well understood. Here, we unravel the neural basis of one such critical computation, namely, location-invariance. This property permits spatial selection to operate no matter which specific locations in the sensory world the competing stimuli occupy. Although appearing straightforward, the implementation of location-invariant stimulus selection requires comparisons between all possible pairs of stimulus locations and is computationally complex: the number of location-pairs at which two competing stimuli could be placed, *L*^2^*-L/2*, scales quadratically with *L*, the number of spatial locations that are encoded. How does the brain meet the resulting demands imposed on neural circuitry and solve location-invariant stimulus selection?

A brain network with a well-established role in spatial target selection, and therefore, an excellent locus to study this question, is the midbrain selection network. It includes the sensorimotor hub, the superior colliculus (SC; or the optic tectum, OT, in birds), and a satellite inhibitory nucleus called the lateral tegmental nucleus ^4, 5^, or isthmi pars magnocellularis, Imc, in birds ^6, 7^ (Supplementary Fig. 1a). The SC/OT, which encodes a topographic map of sensory (and motor) space ^8, 9^, plays a critical role in stimulus selection across spatial locations. Specifically, the intermediate and deep layers of the SC (SCid; called OTid in birds) are required for the selection of the highest priority stimulus among distracters independently of the spatial locations of these stimuli ^10, 11^. This location-invariant selection is expressed in the activity of SCid/OTid neurons as response suppression. When one stimulus is presented at any location, the responses of SCid/OTid neurons encoding that stimulus are suppressed by a competing stimulus presented anywhere outside the neurons’ spatial receptive field (RF) ^12–14^. Mechanistically, competitive suppression in the OTid is orchestrated by the GABAergic Imc through its specialized anatomical connectivity with the OT ^6, 15, 16^. Each Imc neuron receives input from a restricted set of neurons in layer 10 of the OT (OT_10_), but projects back broadly across the OTid space map sparing just those neurons that encode the input locations ^6^ (Supplementary Fig. 1b). This anatomy allows the Imc to implement a spatial inverse operation, distributing inhibition to all competing locations in the OTid space map (Supplementary Fig. 1c). The strength of competitive inhibition depends on the priority of the stimulus ^12–14^, and, notably, inactivation of the Imc abolishes this competitive inhibition as well as spatial selection in the OTid ^15, 16^.

In this context, a conceptually straightforward strategy by which the Imc might achieve location-invariant selection in the OTid is illustrated in Supplementary Fig. 1d – a so-called ‘copy-and-paste’ strategy. Should the spatial RFs of Imc neurons be small, resembling those of the input OT_10_ neurons, then simply repeating the Imc-OT circuit module that solves selection for one pair of locations across all location-pairs, would successfully implement location-invariant stimulus selection. However, the precise nature of the spatial RFs of Imc neurons is not well understood. In fact, the vertically large Imc RFs reported in previous work ^17, 18^ lead to a computational paradox (Supplementary Fig. 1e). Here, we set out to investigate the functional properties of Imc neurons as well as the computations implemented by the Imc-OT network in the barn owl. In doing so, we discovered a combinatorially optimized strategy for location-invariant stimulus selection, one that is supported by unusual encoding of visual space by Imc.

## Spatial RFs of Imc neurons have multiple ‘lobes’

We measured the visuospatial RFs of Imc neurons using extracellular recordings (Methods). Individual Imc units were identified by spikesorting single and multiunit data; only those units deemed to be of ‘high quality’ wee included in the analysis (Methods). We found that individual Imc neurons possessed visual RFs with multiple, distinct firing fields or ‘lobes’ (Fig. 1a-h; Supplementary Fig. 2ab). The number of lobes in each RF was estimated in an unbiased manner using a two-step process (Methods): (i) a nonlinear clustering method^19^ to fit different numbers of clusters to the spatial map of firing rates followed by (ii) a model selection method ^20^ to robustly select the optimal number of clusters in the data (Fig. 1c, g, Supplementary Fig. 2c-f). We found that about two-thirds of Imc neurons had multilobed RFs (80/116; see also Fig. 1l).

**Fig. 1.**
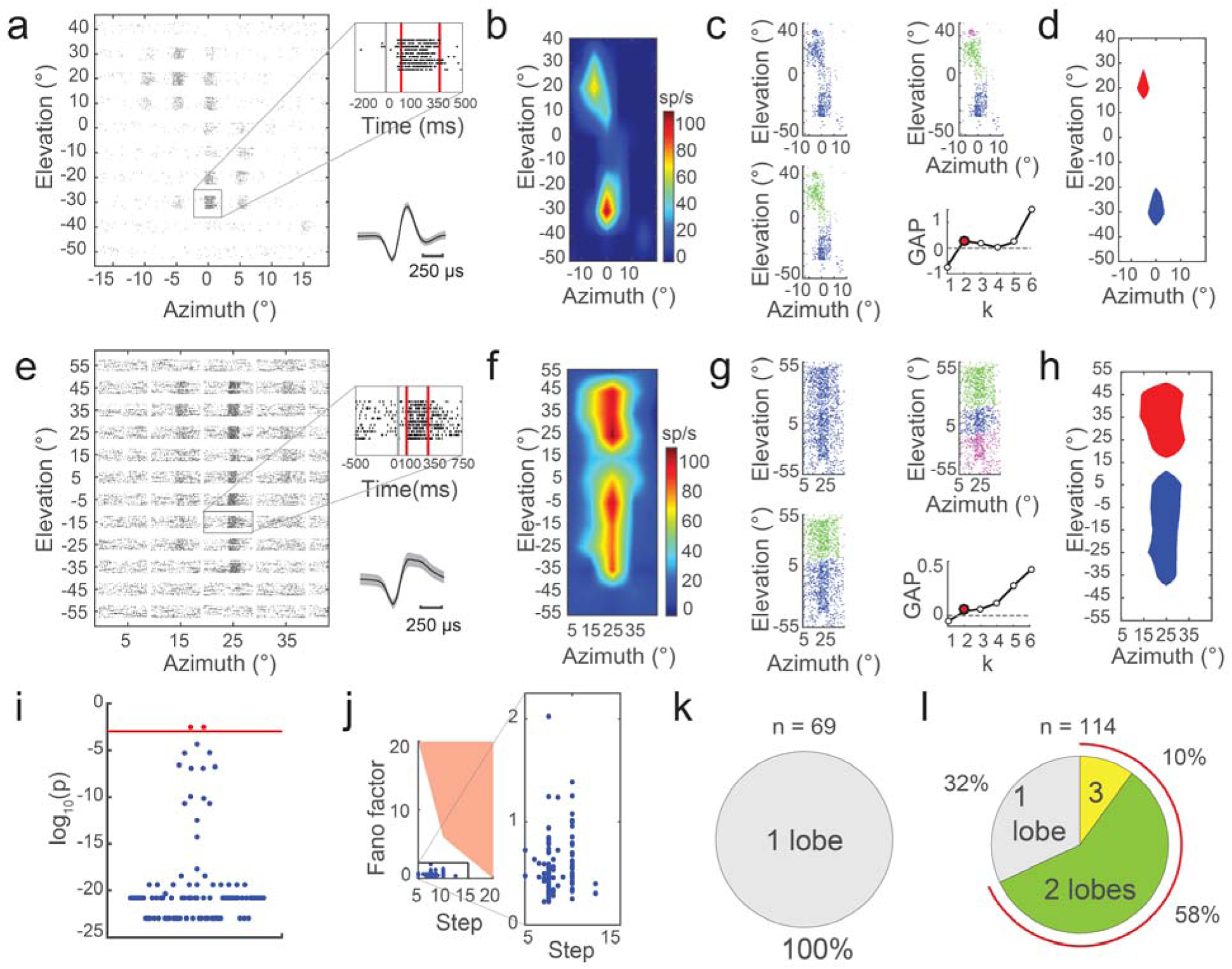
Visual receptive fields (RFs) of Imc neurons have multiple distinct firing fields (‘lobes’). **(a)** 2-D visual RF of Imc neuron: raster plot of neuron’s responses to visual stimulus presented at different spatial locations. *<u>Inset-top</u>*: Gray line – stimulus onset; red lines – time window used to calculate firing rate; evoked firing rates in Imc were high (median = 76.5 Hz; n=114 neurons). *<u>Inset-bottom:</u>* Average spike waveform for neuron in **a**; identified as high-quality unit (Methods); mean (black) ± S.D (gray). **(b)** Color coded firing rate map corresponding to **a**. **(c)** Rate map in **b** re-plotted as distribution of points in a 2-D plane and subjected to spatial clustering (Methods). Shown are the best single (*top-left*), best two (*top-right*), and best three clusters (*bottom-left*) fitted to the data using the density peaks clustering method^19^ (Supplementary Fig. 2c; Methods). *Bottom-right*: Plot of GAP statistic, a robust model selection metric, against the number (*k*) of clusters fitted to data^20^ (Methods). Red point: statistically optimal number of clusters (*k\**), identified as the smallest *k* for which GAP exceeds zero; here *k\** = 2 (Methods) ^20^. **(d)** Half-max extents of these two optimal RF clusters (lobes). **(e-h)** Same as **a**-**d**, but for a different Imc neuron. (**i**) Plot of p-values (logarithmic scale) obtained from separability testing for each sorted unit; one-way ANOVA followed by correction for multiple comparisons (Methods). P-value <0.05 (blue data): units that are deemed ‘well-separated’ from co-recorded units as well as outliers (n=114). Red data: units not well separated form cohort. (**j**) Effect of neuronal response variability and spatial sampling step-size on number of RF lobes detected in a simulated single-lobed Gaussian RF; Monte-Carlo analysis (Supplementary Fig. 2g; Methods). Red area: Fano-factor and step-size pairs yielding >5% rate of misidentifying single-lobed RF as multilobed. Blue data: Experimentally recorded Imc neurons (n = 114). (**k**) Summary of number of RF lobes across 69 OT neurons. See also Supplementary Figs. 1 and 2. **(l)** Summary of number of RF lobes across 114 Imc neurons.

**Fig. 2.**
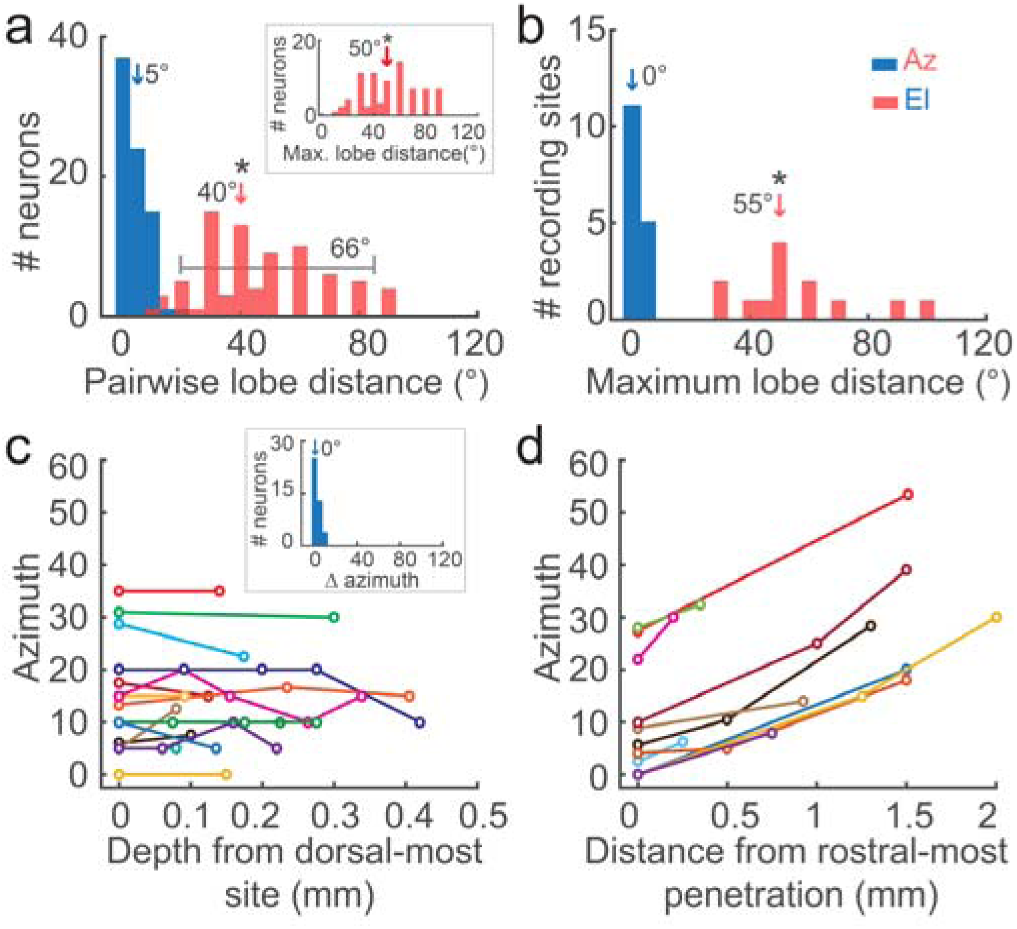
RF lobes of multilobe Imc neurons are distributed along elevation but not azimuth, and RFs are organized topographically in azimuth, but not elevation. **(a)** Histograms of pairwise distances between centers of RF lobes of individual multilobe neurons (Methods). Blue: azimuthal distance; red: elevational distance; marked range: 5^th^ to 95^th^ percentile range of red data. Arrows: median values; *: median significantly different from 0 (p = 0.17, azimuth; p < 0.05, elevation; one-tailed ranksum tests). *Inset*: Histogram of maximum elevational distance between centers of RF lobes of individual multilobe neurons (p < 0.05, one-tailed ranksum test). **(b)** Histograms of maximum distances between centers of RF lobes of multilobe neurons sorted from individual recording sites (Methods); conventions as in **a**; p = 0.65 for azimuth; one-tailed ranskum test, p < 0.05 for elevation; one tailed t-test. **(c)** Plot of average azimuthal center of a recording site against the dorsoventral position of the site within the Imc (Methods); colors: different penetrations. *Inset*: Data re-plotted as histogram of pairwise differences in the azimuthal centers of recording sites along a dorsoventral penetration (p=0.18, one-tailed ranskum test). **(d)** Plot of average azimuthal ‘center’ of a dorsoventral penetration against the rostrocaudal position of electrode in the Imc in that recording session (Methods). Colors: different recording sessions; Spearman correlation =1 in each case. See also Supplementary Fig. 3

To test if the multilobed structure of Imc RFs was an artifact of our experimental methods, we performed three controls. First, we tested if errors in spike sorting might have caused multiple units with single lobed RFs to be misidentified as a single unit with a multilobed RF. To this end, we applied an additional separability criterion to our sorted units. We tested the statistical separability of the waveforms of each sorted unit with those of any other unit as well as with outlier waveforms recorded at the same site, and retained only those units that were well-separated (Methods). We found that the majority of the sorted units (114/116) satisfied the separability criterion as well (p<0.05; Fig 1i), ruling out multiunit contamination as a source of error. Second, we examined if the spatial sampling resolution used for RF measurement, as well as neuronal response variability, might have caused the erroneous identification of single-lobed RFs as being multilobed (Supplementary Fig. 2g). Using experimentally grounded simulations, we mapped out the values of sampling step-size and response Fano-factor that yielded a multilobe misidentification rate of 5% or greater (Fig. 1j; red zone^20^; Methods). We found that the values of these parameters for each recorded unit fell outside the 5% misidentification zone. As a final control, because it is well established that OT RFs have single spatial firing fields, we measured visual RFs of OT neurons. Our methods correctly identified all of the measured OT RFs as being single-lobed (Fig. 1k; Supplementary Fig. 2h). Together, these results confirmed the veracity of our conclusion that the Imc contains predominantly ‘multilobe’ neurons (68%; 78/114; Fig. 1l).

## RF lobes are distributed along the elevation, but not azimuth

To investigate organizing principles underlying spatial encoding by Imc neurons, we analyzed the properties of the measured visual RFs along the two major anatomical axes of the Imc (Supplementary Fig. 1a). The azimuthal centers of RF lobes were nearly identical for lobes within individual multilobe neurons (Fig. 2a, blue data; Methods), across neurons recorded at a given site (Fig. 2b, blue data), and across sites recorded along the dorsoventral axis of the Imc (Fig. 2c; Methods). However, azimuthal encoding varied systematically along the rostrocaudal axis of the Imc: centers of RF lobes encoded progressively more peripheral azimuths as the recording electrode was moved from rostral to caudal portions of the Imc (Fig. 2d ^17, 18^).

The encoding of elevation by Imc neurons was strikingly different. RF lobes of individual multilobe neurons were spaced arbitrarily in elevation (Fig. 2a: large range of red data). Additionally, RF lobes of multilobe Imc neurons were distributed widely across elevational space: for each multilobe neuron (Fig. 2a, inset: large median of data), across neurons recorded at a given site (Fig. 2b, red), and across sites recorded along both dorsoventral and rostrocaudal axes (Supplementary Fig. 3a-d). There was also no systematic relationship between encoded elevations and distance along either principal axis (Supplementary Fig. 3ab).

**Fig. 3.**
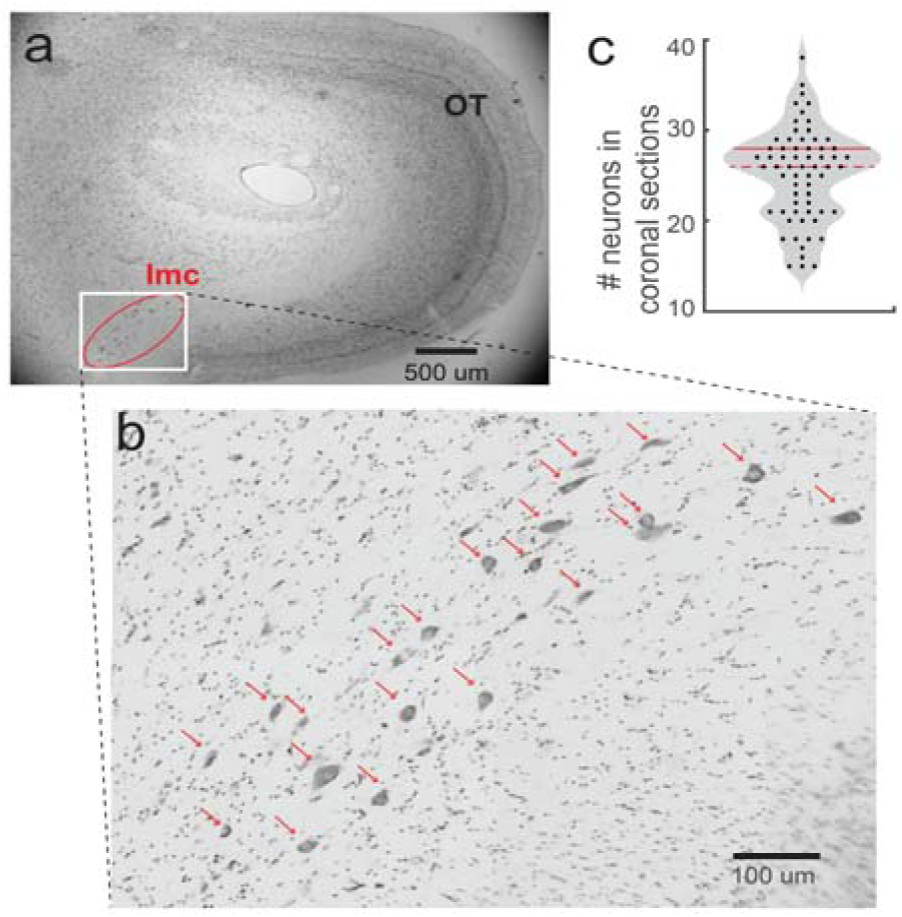
Imc encodes elevations with a sparse number of neurons. **(a)** Coronal section of owl midbrain showing Imc and OT. **(b)** Zoomed-in image showing individual, Nissl-stained, Imc somata (arrowheads); 23 somata in this section. **(c)** violin plot showing number of Imc somata per coronal section; each dot – one section; n=64 sections across two owls. Dashed line: median (26 neurons); solid line: 75^th^ percentile (28 neurons).

These results demonstrated that whereas azimuthal space is encoded in a topographic manner along the rostrocaudal extent of the Imc, elevational space is encoded by multiple, arbitrarily spaced, and widely distributed lobes of varying number and size (Supplementary Fig. 3e-j), with a maximum of three RF lobes per neuron (Fig. 1i).

## Neuronal scarcity necessitates multiple RF lobes

The multilobed encoding of (elevational) space by Imc neurons was puzzling. This was especially so because neurons that provide input to the Imc (OT_10_), as well those that receive Imc’s output (OTid), all tile sensory space with single-lobed spatial RFs organized topographically in both elevation and azimuth (Fig. 1m) ^8^. Might the implementation of stimulus selection across space, a main function of the Imc ^16^, impose any demands on its spatial coding properties? We turned to theory to examine the implications, specifically, of the need for location-invariant stimulus selection on Imc RF structure (Methods). Briefly, we compared the total number of location-pairs at which selection must occur in the OTid, with the number of location-pairs in the OTid at which selection is achievable by a set of Imc neurons. Since multilobed Imc encoding is restricted along the elevation (Fig. 2ab; Supplementary Fig. 3a-d), we focused on stimulus selection between all possible pairs of elevations at any azimuth. We proved mathematically that if the number of Imc neurons (N) encoding different elevations at a given azimuth is less than the number of distinct elevational locations (L) encoded by the OTid at that azimuth (N<L), then multilobed Imc RFs are necessary for location-invariant stimulus selection (Methods).

To examine the biological applicability of this insight, we estimated L and N in the owl brain. For a given azimuth, the OTid encodes elevations ranging typically from −60° to +60° and does so at a spatial resolution of at least 3° ^8, 12^. Consequently, the number of distinct elevational locations encoded by the OTid at a given azimuth is at least 40 (L_el_ > 40). Next, we estimated N_el_. Because visual azimuth is organized topographically along Imc’s rostrocaudal axis (Fig. 2d), transverse sections of the Imc provide snapshots of Imc tissue encoding all elevations at a given azimuth (Fig. 3ab). We obtained histological sections perpendicular to the rostrocaudal axis of the Imc and performed Nissl staining to visualize cell bodies (Methods). Counts of the number of Nissl-stained somata^21^ showed that the majority of sections (75%) had fewer than 28 neurons per section (N_el_; Fig. 3bc). Thus, N_el_ is typically much smaller than L_el_ (median N_el_ / L_el_ < 26/40 = 0.65). In contrast, along the azimuth, there are at least as many Imc neurons as there are encoded azimuthal locations; N_az_ ≥ L_az_ (Methods).

These results indicated that multilobed encoding in the Imc may be driven by the need for the Imc-OT circuit to achieve location-invariant stimulus selection along elevation in the face of a paucity of Imc neurons encoding elevation (Fig. 3bc).

## Model predicts combinatorially optimized inhibition for location-invariant selection

To explore how an under-complete set of Imc neurons might implement location invariant selection, we turned to computational modeling. We set up stimulus selection across spatial locations as an optimization problem with L locations (elevations at a given azimuth), and N model neurons encoding those elevations (N<L; Supplementary Fig. 4; Methods). We imbued all model neurons with Imc-like spatially inverting connectivity with the OT (Supplementary Figs. 1 and 4). The spatial RFs of these model Imc neurons were represented, for simplicity, using ones and zeros, with ones corresponding to locations inside the RF, and zeros, outside (Fig. 4A; also see Supplementary Fig. 4 for validity of model even when this assumption is relaxed). The goal of the optimization was to identify the spatial RF structures of these N neurons (i.e., the numbers of their RF lobes and their spatial locations), such that when two stimuli of equal priority are placed at any pair of locations, they suppress each other equally. This necessary and sufficient condition for location-invariant selection was captured by a specially constructed cost function whose value decreased as the number of location-pairs at which the above condition was satisfied increased. The cost function took the minimum possible value of *–L(L-1)* if and only if the condition was satisfied at all location-pairs (Methods). Any set of Imc RFs that achieved this minimum value, i.e., that achieved location invariant selection, was called an ‘optimal solution’. To match experimental observations (Fig. 1i), we added the constraint that the maximum number of RF lobes allowed for each neuron (*k_max_*) was three.

**Fig. 4.**
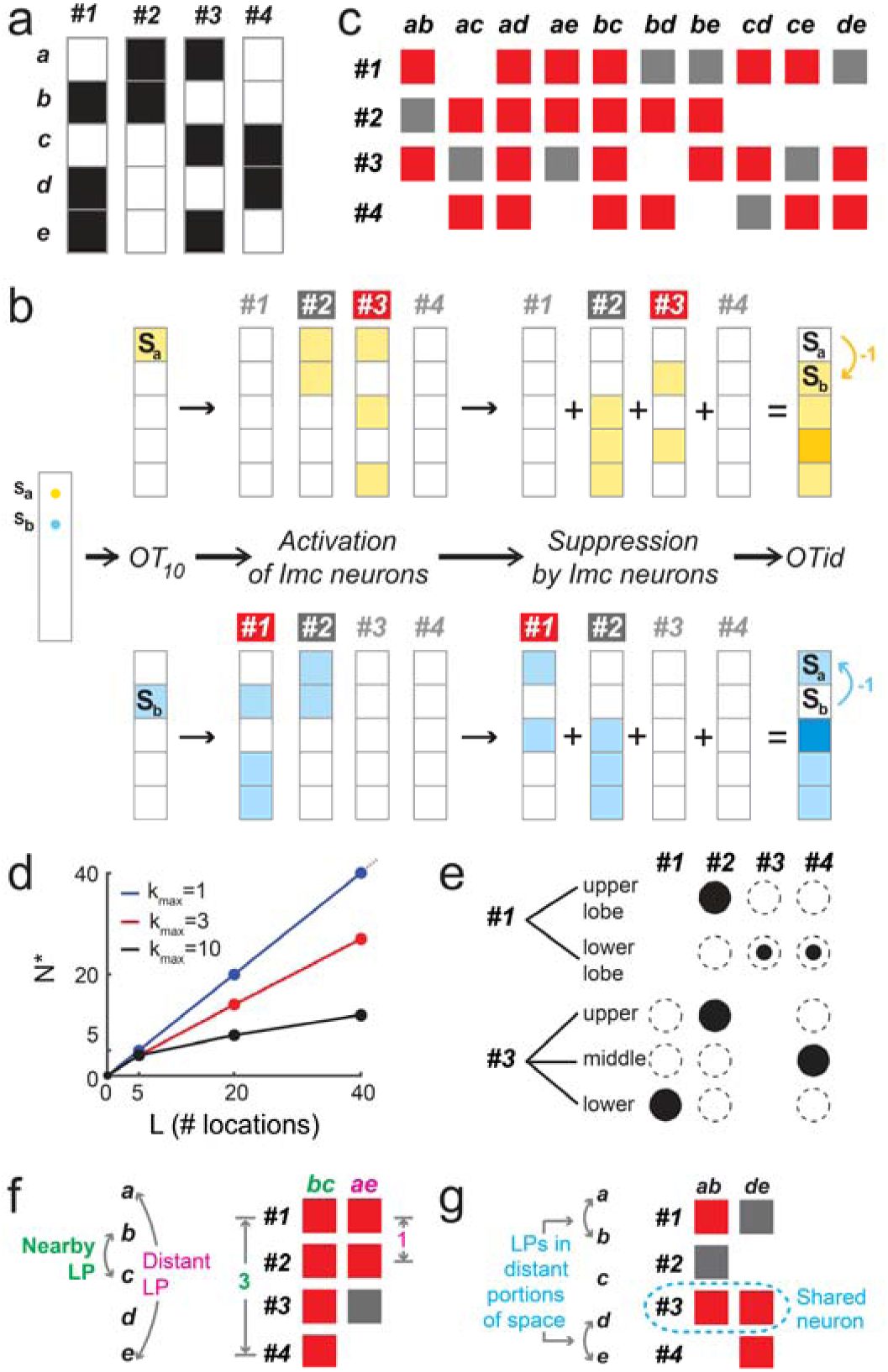
Model predicts combinatorially optimized solution for location-invariant stimulus selection when neurons are scarce. **(a-c)** Illustration of location-invariant selection by an optimal model solution for L=5 locations (*a-e)* and N=4 neurons (*#1-#4)*. (a) The four RFs in the optimal solution. Shaded areas: RF of neuron; two neurons have multilobed RFs *(#1* – two lobes, *#3* – 3 lobes). (b) Optimal solution in **a** implements selection between stimuli S_a_ and S_b_ at location-pair *ab* (*extreme left*). S_a_ and S_b_ are of equal priority (1 unit for simplicity). *<u>Top row</u>:* Information flow through the model OT_10_-Imc-OTid circuit triggered by S_a_. *1^st^ column*: Activation of OT_10_ space map. *2^nd^ column*: Activation of individual Imc neurons. *3^rd^ column*: Suppression pattern generated by each activated Imc neuron (spatial inverse of the neuron’s RF; consistent with published anatomical results; Supplementary Fig. 1b-e ^6^). *4^th^ column*: Combined pattern of suppression in the OTid. Dark colors: 2 units of suppression; light colors: 1 unit (Methods). Curved arrow: Net suppression driven by S_a_ location b. Dark-gray shading: ‘Activated’ neuron (#2); defined as a neuron driven by S_a_ but that does not send inhibition to location b. Red shading: ‘Recruited’ neuron (#3); defined as activated neuron that sends inhibition to location b, thereby involved in selection for location-pair *ab*. *<u>Bottom row</u>*: Same as top row, but for stimulus S_b_. (c) Selection matrix summarizing implementation of selection for all location pairs by optimal model solution in **a**. Columns: 10 possible location-pairs; rows: the four neurons. In each column: dark-gray – activated neurons, red – recruited neurons, blank – neurons not activated by either stimulus. **(d)** Summary plot showing the fewest number of neurons (N*) needed by model to solve location-invariant selection for different numbers of locations (L) (Supplementary Fig. 5c; Methods). *k_max_*: maximum number of RF lobes allowed for each neuron (Methods). **(e-g)** Illustration of signature properties for combinatorially optimized inhibition exhibited by optimal model solution in **a**. (e) Signature property #2 (optimized lobe-overlap; see text). *<u>Top row</u>*: multilobe neuron *#1* in A shares upper, but not lower lobe with neuron *#2*, and shares lower, but not upper lobe with neurons *#3* and *#4*. *<u>Bottom row</u>*: Similar, but for multilobe neuron *#3* (see also Supplementary Fig. 5e). (f, g) Signature property #3 (combinatorial inhibition; see text). *Left panels*: Locations *a-e*. *Right panels*: Patterns of neurons activated and recruited to solve selection for indicated location-pairs (LPs); extracted from selection matrix in **c**. ‘Assortedness’ feature: location-pair *bc* involves nearby locations (f, left panel), but recruits distant neurons to solve selection (f, right panel; *#1* and *#4*; distance =3; Methods); conversely, distant location-pair *ae* recruits nearby neurons (*#1* and *#2*; distance =1. This holds across all permutations of neuronal ordering (Supplementary Fig. 5gh; Methods). Extensive intersection feature: location-pairs occupying distant portions of space (g, left panel) recruit intersecting neural subsets to solve selection (g, right panel; see also Supplementary Fig. 5i; Methods). See also Supplementary Fig. 4, 5 and 6.

An optimal solution for L=5 locations with N=4 neurons illustrates how fewer than L inhibitory neurons can successfully achieve location-invariant selection (Fig. 4a-c; see also Supplementary Fig. 5ab for example optimal solutions for L=20 and L=40 locations). Repeated optimization runs (1000 runs) for L=5 locations and N ranging from 1 to 5 indicated that the smallest number of neurons with which location-invariant selection could be achieved by the model, called N*, was 4 (Supplementary Fig. 5c; Methods). Therefore, the maximum ‘savings’ in the number of Imc-like neurons for L=5 locations was 1 (L-N*). Notably, however, as L increased, neuronal savings increased (Fig. 4d), with L=40 requiring N*=27 neurons to solve location-invariant selection (savings of 13 neurons = 32%; Supplementary Fig. 5b). In addition, neuronal savings also increased as a function of the maximum number of RF lobes allowed per neuron (Fig. 4d).

Further examination of optimal model solutions for all runs of all (L, N*, k_max_) values tested revealed three signature properties that held true in every case. First, every optimal solution contained multilobe Imc neurons (Fig. 4a and Supplementary Fig. 5d). Conceptually, this ‘multilobe property’ is necessary because of the paucity of neurons, i.e., the N<L constraint, as demonstrated by theory (Methods). Second, every multilobe neuron in an optimal solution shared each of its lobes, but not all, with another neuron (Fig. 4e and Supplementary Fig. 5e) – a severe constraint on the relative organization of RF lobes across neurons, one that imposes structured non-orthogonality on the RFs. Conceptually, this ‘optimized lobe-overlap property’ is necessary because selection needs to be solved also when two stimuli are placed at the locations encoded by different lobes of an individual multilobe neuron (Supplementary Fig. 5f). Third, neurons in optimal solutions used a combinatorial inhibition strategy to achieve location-invariant stimulus selection: assorted subsets of neurons were selectively recruited to solve stimulus selection for individual location-pairs, with the subsets corresponding to different location-pairs intersecting extensively. The assorted nature of the subsets was evident in the observation that ‘distant’ neurons were recruited to solve selection between even nearby locations, and vice-versa (Fig. 4f) – features that held true across all permutations of the ordering of the neurons in the solution set (Supplementary Fig. 5gh). The extensive intersection feature was evident in the observation that the neural subsets recruited to solve selection even for location-pairs occupying distant portions of space shared common neurons (Fig. 4g; Supplementary Fig. 5i). Conceptually, this ‘combinatorial property’ is a consequence of the RF lobes of individual multilobe neurons being widely distributed and arbitrarily spaced in optimal model solutions (Supplementary Fig. 5j and 6bd): restricting RF lobes to only nearby locations substantially limits the space of available RF configurations, precluding optimal solutions.

Taken together, the model predicted that the solution of location-invariant selection when N < L necessitated combinatorially optimized coding by sparse, multilobe inhibitory neurons (COSMI). In contrast, when N ≥ L, as is the case with Imc’s azimuthal encoding, the model was always able to solve location invariant selection with single-lobed neurons (Fig. 4d, *k_max_*=1, blue data), using the straightforward copy- and-paste strategy (Supplementary Fig. 1d).

## Experimental validation of model predictions in Imc

To examine if the owl Imc might employ a combinatorially optimized strategy for location-invariant selection in elevation, we tested experimentally whether the RFs of Imc neurons exhibited the three signature properties predicted by the model. Because all elevations at a given azimuth are encoded by neurons within a coronal plane (Fig. 2bc), we sampled these neurons by making recordings at multiple dorsoventral sites within each coronal plane (Methods).

Across recordings made in 16 such coronal planes, we found that multilobe neurons were present in nearly every case (14/16; Fig. 5a; also Supplementary Fig. 3a), thereby validating the signature property #1. The impracticability of recording exhaustively from all Imc neurons in a coronal plane made it infeasible to test if *every lobe* of each multilobe neuron satisfied the optimized lobe-overlap property (signature property #2; Fig. 4e). Therefore, we tested if *at least one lobe* of each multilobe neuron satisfied it (Fig. 5b; Methods). The median fraction of multilobe neurons in each coronal plane that satisfied this property was 1 (Fig. 5c). Finally, we tested signature property #3 (combinatorial inhibition). Both its features, namely, assorted recruitment and extensive intersection, were satisfied in nearly every testable case (7/8 and 6/6 planes respectively; Fig. 5d-g; Methods), despite the non-exhaustive sampling of Imc neurons in individual planes. In addition, the arbitrarily spaced and widely distributed nature of the RF lobes of individual model neurons (Supplementary Fig. 5jk), a model feature driving combinatorial inhibition, was also found in experimental data (Fig. 2ab, Supplementary Fig. 3a-d).

**Fig. 5.**
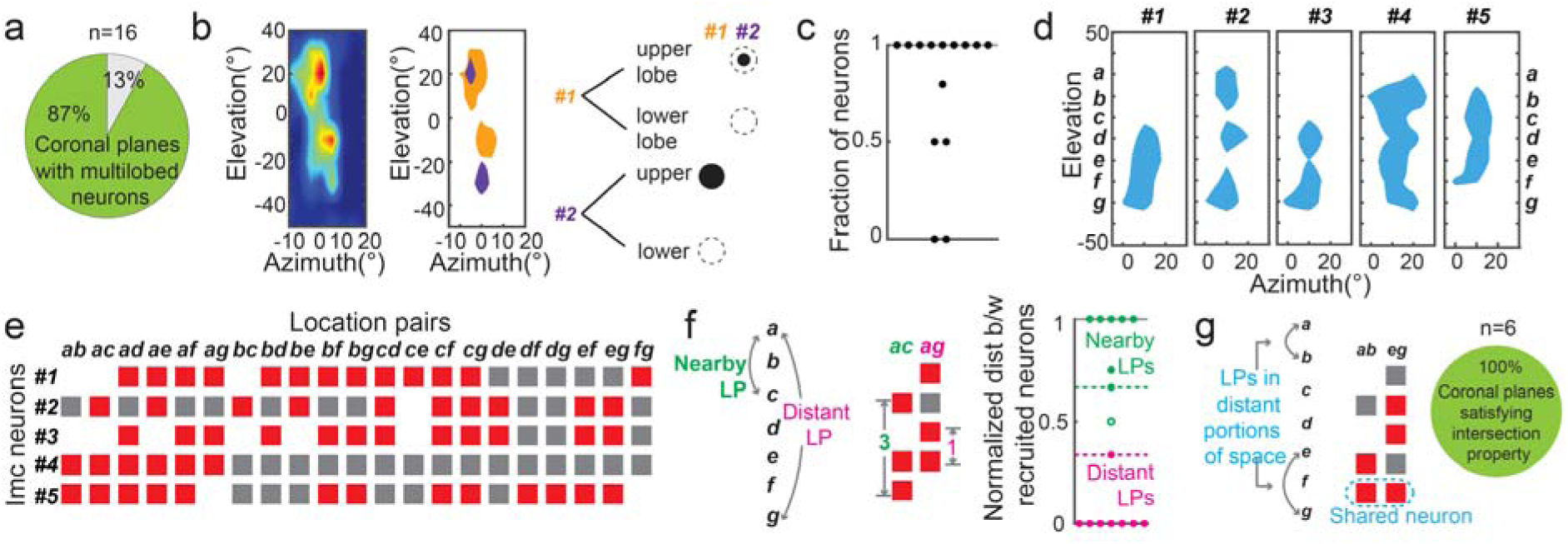
Experimental validation of model predictions in the Imc. (a) Signature property #1: Pie-chart summary of fraction of Imc coronal planes tested that contained multilobe neurons (87% = 14/16 planes; also Supplementary Fig. 3ef). (b-c) Signature property #2. (b) *Left:* Rate map of RF of another Imc neuron sorted from the same recording site as the neuron in Fig. 1a-d. (Only these two neurons were recorded in this Imc coronal plane.) *Middle*: Half-max of RFs of neurons in Fig. 1 (purple; reproduced from 1d) and Fig. 5b left (orange). *Right*: For each neuron, the upper RF lobe, but not lower one, shows overlap, satisfying the testable lobe-overlap property (see text); conventions as in Fig. 4e. (c) Fraction of multilobe neurons in each coronal plane satisfying the testable version of lobe-overlap property; dot – coronal plane; median fraction = 1. (d-f) Signature property #3. (d) RFs (half-max) of all Imc neurons recorded within an example coronal plane. *a-g* are seven (discretized) spatial locations encoded by these neurons (Methods). (e) Selection matrix showing combinatorial activation of recorded neurons for selection at different location-pairs; conventions as in Fig. 4c. (f) *<u>Two left panels</u>*: Illustration of assortedness feature for example in d; conventions as in Fig. 4f (Methods). *<u>Right</u>*: Summary of this feature across Imc coronal planes; only those planes containing ≥ 3 Imc neurons each were testable (8/14; Methods) Dashed lines: Distance cut-offs for ‘distant’ neurons (green; 0.66) and ‘nearby’ neurons (magenta; 0.33; Methods). Filled circles: Imc coronal planes that satisfied these cut-off criteria; ≥ 7/8 in ≥ each case (Methods). (g) *Left:* Illustration of ‘extensive intersection’ feature for example in d; conventions as in 4g. *Right*: Pie-chart summary of this feature across coronal planes (100% exhibited the feature; 6/6). Note that this feature was testable only for those planes for which the recorded neurons encoded location-pairs occupying distant portions of space (6/14; Methods).

## Metabolic and wiring costs explain specialized properties of Imc neurons

Three questions regarding the biological implementation of location-invariant selection in the Imc circuit remained puzzling. First, why might N<L be biologically desirable in the Imc, in the first place (necessitating combinatorially optimized inhibition)? Second, if N<L is attractive biologically, why don’t Imc RFs have a large number of lobes, thereby achieving greater savings in the number of Imc neurons (Fig. 4d)? In other words, why is the maximum number of Imc RF lobes restricted to a low number (k_max_ = 3; Fig. 1i)? Third, why is multilobed encoding found only along one spatial axis (here, elevation), why not along both axes for greater neuronal savings?

To gain insight into these questions, we examined Imc function in the context of two types of costs that nervous systems must incur in building and operating a neural circuit: wiring cost and metabolic cost. We estimated wiring cost by quantifying the cost of implementing spatially inverting projection patterns from the Imc to the OT (Methods ^22^), and metabolic cost by quantifying the cost of broadcasting of spikes across the OT for competitive suppression. We found that wiring cost decreases as the number of RF lobes increases (Fig. 6a; Methods). In contrast, metabolic cost increases as the number of RF lobes increases (Fig. 6b; Methods). Consequently, the wiring cost places a lower bound on the number of RF lobes (and a corresponding upper bound on the number of neurons), whereas the metabolic cost places an upper bound on the number of RF lobes (and a lower bound on the number of neurons). The ideal number of RF lobes (and the number of neurons necessary), therefore, is one that minimizes some weighted combination of the two opposing costs (Fig. 6c; blue). Because Imc neurons have high firing rates (median = 76.5 Hz ^15, 23^; Fig. 1a), the metabolic cost of Imc function scales up substantially, pulling the ideal number of RF lobes to even lower values (Fig. 6c, red vs. blue; thereby also providing a rationale for the continued presence of some single-lobe neurons in the Imc; Fig. 1i)

**Fig. 6.**
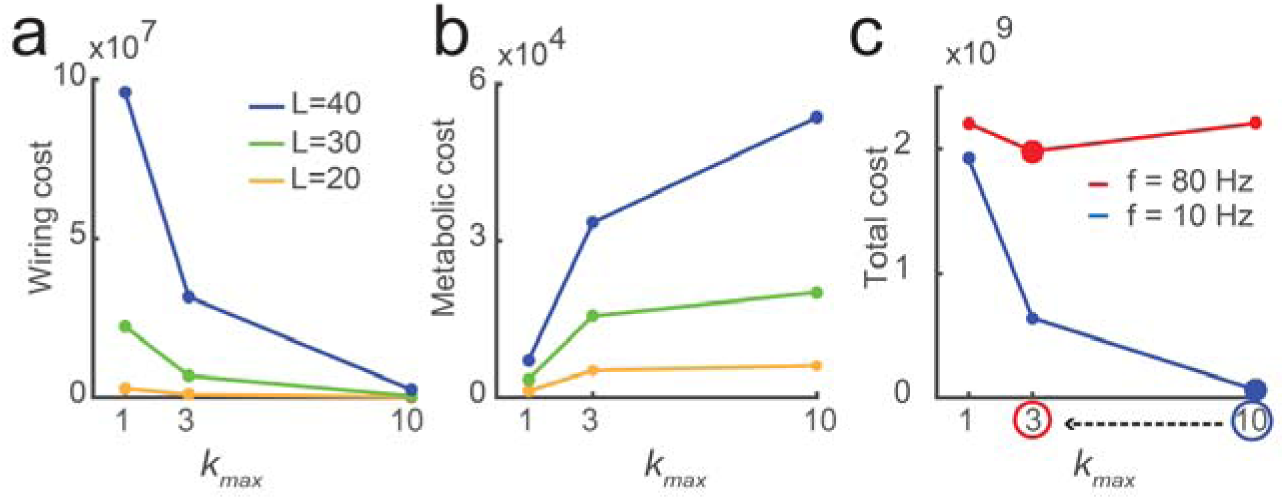
Metabolic and wiring costs of location-invariant stimulus selection. **(a)** Wiring cost plotted as a function of the maximum number of Imc RF lobes allowed (*k_max_*); calculated across optimal model solutions (Methods). **(b)** Metabolic cost as a function of *k_max_* (Methods). **(c)** Schematic showing total cost (weighted combination of **a** and **b**) for Imc circuit to solve location-invariant selection for a given L at low average firing rates (blue: 10 Hz), and high average firing rates (red: 80 Hz; Methods). Circled values along x-axis (and corresponding large dots) indicate the optimal *k_max_* values at the two firing rate levels. Results demonstrate left shift of optimal *k_max_* with increasing firing rates (Methods). Absolute values of optimal *k_max_* are a result of the specific weights chosen here; weights identical for both curves. In all cases: mean ± SD values are plotted; SD values smaller than size of dots.

Taken together, these results indicate that a small number of Imc neurons (N<L), with multilobed RFs that have a small number of RF lobes (small k_max_ value), are ideally suited to achieve location-invariant selection while minimizing the net neural costs. Therefore, increasing excessively the number of RF lobes along one spatial axis (here, elevation), or increasing the number of RF lobes also along the other axis as well (here, azimuth), is not biologically desirable. The occurrence of multilobed encoding specifically along elevation, rather than azimuth, is likely a side-effect of azimuthal inputs from OT’s rostrocaudal axis being mapped directly onto the parallel (and long) rostrocaudal axis of the Imc ^6^, relegating elevation to be coded by the transverse (and neurally sparse) planes.

## Discussion

The combination of electrophysiology, theory, anatomy, and modeling in this study provides a detailed unpacking in owls of a critical neural function, namely location-invariant stimulus selection.

### Multilobed visuospatial RFs and stimulus selection

Multilobed spatial RFs have not been reported previously in any visual sensory area to the best of our knowledge. We find that in the Imc, a sensory area that is just two synapses away from the retina ^24^, the majority of neurons have multilobed visual RFs. This contrasts with previous reports of large, vertically elongated visual RFs in the Imc ^17, 18^ (a consequence of the detailed approaches used here, rather than species differences ^15^). Multilobed Imc RFs were uncovered here using flashing dots as visual stimuli, a classical approach that has been used extensively in visual neuroscience studies across species. The use of this approach contrasts directly the unusual multilobed encoding of space by Imc with the single-lobed encoding of space by OT (Fig. 1k).

We demonstrate the need for such unusual encoding in the inhibitory Imc (Fig. 3), and uncover a novel neural strategy for location-invariant stimulus selection – combinatorially optimized feature coding by a sparse set of multilobe inhibitory neurons (COSMI; Fig 4). The need for this strategy is unimpacted by the simplifying assumption of binary RFs made by the optimization model (Supplementary Fig 4), and is further supported by experimental validation of model predictions (Fig. 5). Additionally, through subsequent estimation of the net cost of neural circuit operation, we provide a plausible rationale for ‘why’ the owl Imc may be organized, anatomically and functionally, in the way that it is (Fig. 6). The specific values of k_max_, the maximum number of RF lobes, used to develop this rationale (Fig. 6) represent values that are particularly relevant to the Imc: k_max_=1 corresponds the single-lobed case, k_max_=3 corresponds to the experimentally determined value in the owl Imc, and k_max_=10 corresponds to the practical upper bound on the number of possible RF lobes (based on the functional properties of Imc neurons; Methods).

The arguments in this study are framed in the context of selection between pairs of locations. Because selection among multiple stimuli requires comparisons between all possible pairs of stimulus locations, the computational principles uncovered here apply directly to the general problem of selection across an arbitrary number of competing stimuli.

### COSMI is distinct from traditional population coding schemes

Combinatorially optimized coding is conceptually distinct from traditional population neural coding schemes. For instance, in population vector coding, multiple neurons with overlapping, single-lobed tuning curves (or RFs) are activated to encode feature values such as stimulus locations, motion direction, etc., with high precision ^25–28^. Typically, it is possible to order these RFs along the feature axis such that neighboring values of features are always encoded by functionally ‘local’ subsets of neurons (Supplementary Fig. 6ac). In contrast, neurons with multilobed RFs cannot be ordered this way: some neurons always code also for distant locations (Supplementary Figs. 6bd and 5i), and selection for a given location-pair cannot be guaranteed by a ‘local’ subset of neurons (Supplementary Fig. 6bd). A population coding scheme reported in the literature that does involve multilobed encoding as well as the activation of non-local neural subsets is the combinatorial coding of odors by olfactory receptor neurons ^29^. Whereas assorted and extensively intersecting subsets of neurons are activated to encode odors, no inherent constraint on the relative positioning of these RF lobes across neurons has been demonstrated. In contrast, in the combinatorially optimized coding reported here, the placement of RF lobes needs to be optimized across neurons, and is exemplified by the lobe-overlap property (Fig. 4e). For this same reason, our scheme also differs from the encoding of space by entorhinal grid cells: the firing fields of different grid cells are not inherently yoked to one another ^30, 31^. In addition, each grid cell has a large number of highly organized firing fields, unlike the few, and arbitrarily placed, RF lobes of Imc neurons. Finally, combinatorially optimized coding also stands in direct contrast to the sparse, orthogonal coding by an overcomplete set of neurons reported in many brain areas ^32, 33^: it involves promiscuous, non-orthogonal coding by an under-complete set of neurons. The problem of location-invariant selection with limited neurons, which yields combinatorially optimized coding in Imc, belongs to the same (np-complete) class of computationally complex problems as the traveling salesman problem and the minimum spanning tree problem ^34, 35^. Although the brain solves it naturally, exactly how Imc’s optimized, multilobed RFs are specified during neural development is an intriguing open question.

### Generality of COSMI beyond the owl Imc

The discoveries, here, of multilobed visual representation, a new form of population coding, and an efficient neural solution for a critical brain function (stimulus selection) have come from the systematic study of the functional response properties of inhibitory neurons in the owl Imc.

The Imc, called the periparabigeminal lateral tegmental nucleus (pLTN) in mammals, is conserved across vertebrate midbrains, as is the specialized anatomical connectivity between the Imc/pLTN and the OT/SC ^5, 6^. It is the primary source of long-range competitive inhibition to the OT ^16^, and has been proposed to be a critical processing hub for stimulus selection for attention ^7, 15, 16^. However, the functional properties of this midbrain nucleus of emerging importance have not been studied in any vertebrate other than the barn owl thus far. The biological advantages afforded by combinatorially optimized inhibition together with the Imc’s conserved nature suggest that COSMI may be a solution employed generally by the vertebrate midbrain to achieve location-invariant spatial selection.

The computational principle of combinatorially optimized inhibition also extends naturally to selection across values of other stimulus features such as orientation, color, odor, etc. Typically, the functional properties of inhibitory neurons in cortical as well as sub-cortical areas are less well-studied than those of primary (pyramidal) neurons. Our results indicate that a careful examination of the encoding properties of inhibitory neurons in key brain areas may reveal COSMI, the result of concerted shaping of functional and structural circuit properties, as a widespread strategy for efficient, feature-invariant stimulus selection and decision-making under metabolic and anatomic constraints.

## Acknowledgements

This work was supported in part by funding from the Science of Learning Institute (JHU), and NIH R01 EY027718. We are grateful to Drs. James Knierim and Daniel O’Connor for feedback.

## Author Contributions

NRM and SPM designed the research, performed experiments, analyzed the data and wrote the paper.

## Methods

### Animals

We performed experimental recordings in 15 head-fixed, non-anesthetized adult barn owls that were viewing a visual screen passively (*Tyto alba)*. Both male and female birds were used; the birds were shared across several studies. All procedures for animal care and use were carried out following approval by the Johns Hopkins University Institutional Animal Care and Use Committee, and in accordance with NIH guidelines for the care and use of laboratory animals. Owls were group housed in enclosures within the aviary, each containing up to 6 birds. The light/dark cycle was 12 hrs/12 hrs.

### Neurophysiology

Experiments were performed following protocols that have been described previously ^12, 16^. Briefly, epoxy-coated, high impedance, tungsten microelectrodes (A-M Systems, 250µm, 5 −10 MΩ at 1 kHz) were used to record single and multi-units extracellularly. A mixture of isoflurane (1.5-2%) and nitrous oxide/oxygen (45:55 by volume) was used at the start of the experiment to anesthetize the bird and secure it in the experimental rig (a 30-minute period of initial set-up). Isoflurane was turned off immediately after the bird was secured and was not turned back on for the remainder of the experiment. Frequently, nitrous oxide was also turned off at this point, but in several experiments, it was left on for a few hours if the bird’s temperament necessitated it (some birds were calm when restrained, while others were not). However, it was turned off at least 30 minutes before the recording session. Our recordings were performed starting, typically, 3 hours after initial set-up (the time required for positioning the electrode). As recovery from isofluorane occurs well under 30 minutes after it is turned off, and recovery from nitrous oxide occurs within a minute (the bird stands up and flies away if freed from restraints), recordings were made in animals that were not anesthetized and non-tranquilized.

We first targeted the OT based on well-established methods ^8^. We then navigated to the Imc using the OT’s topographic space map as reference and previously published methods (Supplementary Fig. 1a) ^16^. The Imc is located approximately 16 mm ventral to the surface of the brain. Dorsoventral penetrations through the Imc were made at a medial-leading angle of 5° from the vertical to avoid a major blood vessel in the path to the Imc.

### Visual stimuli and RF measurement

Visual stimuli used here have been described previously ^12, 13^. Briefly, either stationary, translating, or looming visual dots (of fixed contrast) were flashed at different locations on a tangent TV monitor in front of the owl. Looming stimuli were dots that expanded linearly in size over time, starting from a size of 0.6° in radius. Visual stimuli were presented for a duration of 250ms (and inter stimulus interval of 1.5-3 s) at all sampled locations. Pilot experiments indicated that visual RFs were narrow in azimuth but spread along the elevation. Therefore, RF measurements were made by presenting stimuli over the −60° to 60° range in elevation, and over a 40° (± 10.4°) range in azimuth (centered around the azimuth that yielded the best responses). Each sampled stimulus location was repeatedly tested 9-15 times in a randomly interleaved fashion. Multi-unit spike waveforms, recorded using Tucker Davis Technologies hardware interfaced with MATLAB, were sorted off-line into putative single neurons (see below). The spatial responses for each neuron were measured by counting spikes at each sampled location during a 100-350 ms time window following stimulus onset.

### Spike sorting multi-unit data

The ‘*chronux’* spike-sorting toolbox was used for the majority of the analyses ^36^. This method is based on a hierarchical unsupervised clustering approach in which the spike waveforms are initially classified into a large number of clusters, typically 10 times the number of putative units recorded. Clusters with very few spikes are discarded and the remaining clusters are then aggregated automatically using metrics of similarity between waveform shapes. In addition, we include only those units for analysis that have less than 5% of the spikes within 1.5 ms of each other (ISI criterion).

The statistical separability of individual sorted units was assessed based on the distance of a unit’s cluster (of waveforms) from the clusters corresponding to other units as well as the outlier cluster measured at the same site. We first projected the spike waveforms measured at a given site to a 3-dimensional space using principal components analysis. Then, we performed a one-way ANOVA test to examine if the mean of the waveforms of a given unit (in the projected 3-dimensions) was significantly different from the means corresponding to the other units and the outliers. This was followed by the Holm-Bonferroni criterion for multiple comparisons. In a few cases (4/116), there were either too few waveforms in the outlier cluster (number of waveforms in outlier cluster < 8% of number of waveforms in any of the remaining sorted units), or the outlier waveforms did not form a cluster with a Gaussian distribution. In such cases, we only tested for the distance of the unit’s cluster mean from the cluster means of other units. We regarded only those units whose cluster means were significantly different from the means of all other units (and the outlier cluster) as ‘well-separated’ units per this separability criterion (p<0.05; the p-value plotted for each unit in Figure 1i is the largest p-value obtained across all comparisons for that unit). Only well-separated units were included in all remaining analyses (subsequent to Fig. 1i) in this study.

### Identification of the optimal number of RF lobes (Fig. 1)

In order to determine the number of firing fields (or lobes) in an RF in an unbiased manner, we first transformed the measured RF responses to a distribution of points in 2-dimensional space (azimuth x elevation). This distribution was generated such that the density of points around each sampled spatial location was proportional to the firing rate of the neuron evoked by a visual stimulus presented at that location. We achieved this by distributing points randomly and uniformly within a rectangle centered around the sampled location such that the number of points was equal to the firing rate at that location; the dimensions of the rectangle were the azimuthal and elevational sampling steps, respectively. This transformation allowed us to apply spatial clustering methods to the firing rate maps.

Next, using the density peaks clustering method ^19^, we fit successively *k*=1,2,3…6 clusters to the distribution (Fig. 1cg). This clustering method identifies cluster centers by searching for regions that have high local density of points (ρ) that are also far away from any points of equal or higher density (δ=minimum distance from points of equal or higher density; Supplementary Fig. 2c-f. For the point with highest local density, δ is conventionally taken as the maximum distance of the point from all other points). It is robust to nonlinear cluster boundaries and unequal cluster sizes – conditions under which traditional methods like k-means perform poorly. The *k* cluster centers are chosen by the algorithm as points with the *k* highest values of gamma (*γ*), defined as the product of ρ and δ. We repeated this procedure for each k, thereby fitting the 1-best, 2-best, … 6-best clusters to the data.

Following this, we applied a model selection procedure to identify the optimal number of clusters in the data, i.e., the best *k* value (*k\**), based on the ‘gap statistic’ ^20^. This is an unbiased method to detect the number of clusters that best fit a distribution of points. For each k, we estimated a ‘gap’ value (*gap(k)*), which evaluated the goodness of fitting k clusters to the distribution. The gap value was calculated by standardizing the pooled within-cluster sum of square distances between all points in each of the k clusters (*W_k_*) and comparing its log value (log (W_k_)) to the expectation of this quantity, (E*(log (W_k_)), under the null hypothesis that the data contains only one cluster ^20^. We calculated this in MATLAB by using the *‘evalclusters’* function with *‘gap’* as the evaluation method, which yielded *gap(k*) as well as *se*(k) for each k; *se(k)* was the standard error in the estimate of *gap(k).* Then, the gap selection statistic was defined as, *GAP(k) = gap(k)-gap(k+1) + se(k+1)*. The number of clusters that fit the data optimally is defined by the method as the smallest value of k for which *GAP(k)* >= 0. Conceptually, the value of *GAP(k)* for the null hypothesis (*k\**=1) keeps decreasing linearly with increasing *k*, whereas the rate of the decrease of the metric under the alternate hypothesis (*k\**>1) has been shown to fall exactly at *k=k*.* Hence the *‘gap’* between the two curves is maximum at *k=k*,* and *GAP(k)*, the difference between *gap(k)* and *gap(k+1)* is greater than zero for the first time when *k=k\**.

### Defining the centers of RF lobes

The center of an RF lobe defined as the stimulus location evoking the highest firing rate within that lobe. The azimuthal RF ‘center’ of an Imc neuron is defined as the average of the azimuthal centers of all of its RF lobes, because RF lobes of an individual neuron do not vary significantly in azimuth (Fig. 2a; blue). The azimuthal RF ‘center’ of a recording site in the Imc, across all the neurons recorded at that site, is defined as the average of the azimuthal centers across all the RF lobes of all the neurons recorded at that site. This is valid because RF centers of individual neurons within a recording site do not vary significantly in azimuth (Fig. 2b; blue). The azimuthal RF ‘center’ of a penetration is defined as the average of the azimuthal centers across all recording sites in that penetration. This is valid because RF centers of individual recording sites within a penetration do not vary significantly in azimuth (Fig. 2c).

### Monte-Carlo analysis of the effect of neuronal noise and spatial sampling resolution on number of detected RF lobes (Fig. 1)

A low spatial sampling resolution during the measurement of spatial RFs, as well as high variability in neural responses, could both cause a single lobed RF to appear falsely as a multilobed one (see Supplementary Fig. 2g). To test how robust our method for identifying the ideal number of RF lobes is to sampling resolution (sampling step-size) and neural response variability (response Fano-factor; defined as variance/mean), we performed the following control. First, we generated a single-lobed Gaussian in 2D (azimuth x elevation), with mean and covariance equal to the average values of these parameters across all the experimentally measured Imc RFs (114 Imc units). Using this single-lobed Gaussian as ‘reference’, we repeatedly simulated RFs using different step-sizes and different response Fano-factor values: For a given step-size, the firing rate at each location was chosen randomly from a normal distribution with mean equal to the value yielded by the reference Gaussian at that location, and variance determined by the Fano-factor value. Next, we transformed this simulated RF into a distribution of 2-D points and applied the density peaks clustering method. Finally, we applied the gap-statistic model selection method to determine the ideal number of lobes in the RF. We repeated this 150 times for each step-size and Fano-factor pair, and calculated the fraction of times for which multiple RF lobes were detected (erroneously) in this data. We repeated the whole procedure for a range of step-size and Fano-factor values that subsumed the range of experimental step-sizes and measured Fano-factor values, and identified the zone that yielded ≥ 5% false detection rate of multiple lobes (Fig. 1j).

To test if our experimental conditions had a high chance of falsely detecting multilobed RFs, we compared the experimentally used step-size for each RF and the RF’s Fano-factor value with those that yielded a ≥ 5% false detection rate in simulation. The Fano-factor for each RF was calculated as the average of the Fano-factor values at all sampled locations in that RF. The step-size for each RF was calculated as the average of the azimuth and elevation sampling steps used to measure the RF. We found that all of our RFs were well within the ‘safe’ zone of ≤ 5% error (Fig. 1j). Thus, the detection of multilobed RFs in our data was unlikely to be a spurious consequence of sub-optimal measurement conditions.

### Theoretical calculations regarding the need for multilobed RFs

We wondered if the implementation of stimulus selection in the OTid, specifically, location-invariant selection in the OTid, imposed any demands on Imc RF structure. To this end, we compared the total number of location-pairs at which selection must occur in the OTid, with the number of location-pairs at which selection is achievable by a set of Imc neurons. Since multilobed Imc encoding is restricted along the elevation (Fig. 2ab), we focused on stimulus selection between all possible pairs of elevations at any azimuth.

#### Simplified version

We started by making two simplifying assumptions: (a) that the OT space map is a collection of non-overlapping spatial RFs that tile sensory space, and (b) that each Imc neuron has exactly *k* RF lobes (*k* always ≥ 1).

In this scheme, if the number of distinct elevations (at a given azimuth) in the discretized OT space map is *L*, then the total number of distinct pairs of stimulus locations possible is *L(L-1)*. A stimulus placed within any RF lobe of a *k*-lobed Imc neuron can suppress competing stimuli located anywhere outside the RF, i.e., at *L-k* locations. Therefore, each Imc neuron is capable of implementing competitive selection at *k(L-k)* pairs of locations. With *N* such Imc neurons, the number of pairs of stimulus locations at which competitive selection can be resolved by the Imc is at most *Nk(L-k)*. Note that this quantity is computed assuming no overlap between Imc RFs and is greater than the number of pairs of stimulus locations at which competitive selection can be resolved by the Imc if overlap between RFs is allowed. Therefore, to achieve successful competitive suppression between all possible pairs of stimulus locations, i.e., location invariance, a condition that must be satisfied is

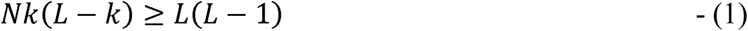

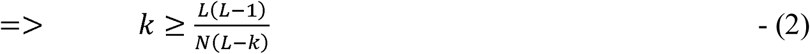

This necessary (but not sufficient) condition for location invariance is already very revealing: If all Imc neurons had only single lobed RFs, i.e., *k* = 1, the above inequality reduces to *N* ≥ *L* i.e., the number of Imc neurons would need to be greater than or equal to the number of distinct spatial locations. Since the logical proposition ‘A => B’ is exactly the same as the proposition ‘not (B) => not (A)’, in our case, the proposition ‘k = 1 => N ≥ L’ is exactly the same as the proposition ‘N < L => k≠1’, i.e., if the number of Imc neurons is less than the number of spatial locations, then at least one Imc RF must be multilobed (because RFs cannot have fewer than one lobe, by definition).

This conclusion held true even when both the simplifying assumptions – (a) that OT RFs are non-overlapping, and (b) that all Imc neurons have the same number of RF lobes – were relaxed (see ‘Full version’ next).

#### Full version

We used a more biologically accurate model of space in which RF extents, overlap of RFs across neurons, and the resolution of competition reported in the OTid (the minimum distance between two stimuli such that OTid is able to select the stronger of the two stimuli) ^12^ were all modeled to match experimental data. In addition, we allowed varying numbers of Imc RF lobes:

Let the total range of elevational locations for which barn owl’s midbrain encodes space be R and the resolution of encoding space be r. Then, the number of distinct locations at which a stimulus can be placed along elevation is 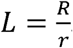 Let the resolution for competitive selection be *Cres*.

The total number of distinct location-pairs at which two competing stimuli can be placed such that they are greater than *C_res_* apart from each other is approximately 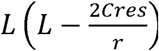. Note that this quantity is calculated by counting all the locations at which a second stimulus can be placed such that it is at least *C_res_* away on either side of a first stimulus that is placed in any of the *L* locations. However, when a first stimulus is placed at the edge of the visual field, a second competing stimulus can be placed only on one side such that it is *C_res_* away. It is straightforward to show that 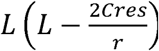 is smaller than the quantity when we include the edge effects. Hence, for location invariance to be achieved, selection of the stronger stimulus must at least be implemented when two competing stimuli are placed in any of these possible location-pairs.

Let the number of lobes in a given Imc neuron be *k* Let the half-max size of each lobe be *l_h_*. Then, a k lobed Imc neuron solves competition for a total of 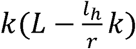 location-pairs (assuming each Imc neuron sends inhibition to all locations that lie outside the half-max extent of the neuron’s RF, without loss of generality; see “Model assumptions” section below and Supplementary Fig. 4 for implications of this assumption). This is just the number of location-pairs such that one stimulus can be placed inside the multi-lobed RF (at its peak for effective suppression of competing stimuli) and the other outside. Let the total number of k lobed Imc neurons be *N_k_*

Therefore, the total number of Imc neurons is

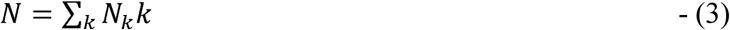

To achieve location invariance, we need

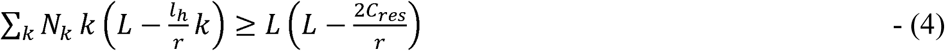

Since *k* ≥ 1, and *l_h >_*_2*cres*_, (mean *l_h_*= 33.6°± 1.25° from the 209 RF lobes across 114 Imc neurons we measured, and C_res_ < 10° ^12^), we get

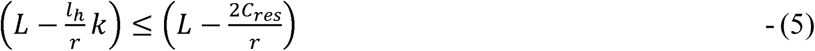

Using (5) in (4) gives,

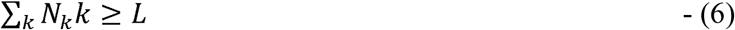

In other words, if all the Imc neurons are single lobed (k=1), this equation becomes *N* ≥ *L*. Since the logical proposition ‘A => B’ is exactly the same as the proposition ‘not (B) => not (A)’, the proposition *‘k = 1 => N L’* is exactly the same as the proposition *‘N < L => k* ≠ *1’* i.e., if the number of Imc neurons is less than the number of spatial locations, then at least one Imc RF must be multilobed (because RFs cannot have fewer than one lobe, by definition).

#### Histology (Fig. 3)

Owls were perfused with paraformaldehyde and their brains extracted per standard procedures. The fixed brains were blocked so that the rostro caudal axis of the Imc was perpendicular to the sectioning plane, and brain sections of 40 µm thickness were obtained. Sections containing Imc were mounted, Nissl stained, and cover-slipped. Sections were imaged at 40x under a light microscope and the number of Nissl stained somata in the Imc in each section were manually counted by NRM and SPM independently^21^. For each section, the maximum value of the counts from the two authors was used to generate the plot in Fig. 3c.

#### Location-invariant selection across azimuthal locations

The OTid encodes azimuths ranging typically from −10° to 60° at a spatial resolution of no better than 1° ^8, 12^. Consequently, the number of distinct azimuthal locations encoded by each OTid is ≤ 70 (L_az_ ≤ 70).

Because the rostrocaudal extent of the Imc is 2800 µm, and the somas of Imc neurons are no larger than ∼33 µm (largest somatic dimension = 33 µm, n=456 neurons across 20 coronal sections), there are at least 70 (coronal) sections along the rostrocaudal axis of the Imc, with each section containing at least one Imc neuron not also found in the neighboring sections.

Consequently, there are at least 70 neurons involved in encoding the L_az_ distinct azimuths, N_az_ ≥ 70; N_az_ ≥ L_az_. (For this conservative estimate of N_az_, we only need that of the ∼26 neurons in each successive coronal section of the Imc (median #neurons per section = 26; Fig. 3c; dashed red line), just one be distinct.

Thus, there are sufficient Imc neurons to encode azimuthal locations, precluding the need for a combinatorial solution for location invariant selection along the azimuth (involving multilobe neurons with RF lobes spread along the azimuth). Consistent with this expectation, azimuthal encoding by Imc neurons is effectively single-lobed: all lobes of a multilobe Imc neuron encode the same azimuth (Fig. 2a-c).

### Optimization model for solving location-invariant stimulus selection across elevations (Fig. 4)

#### Conceptualizing and setting-up the model (Supplementary Fig. 4)

In our model,

- *L* = number distinct spatial elevations at a given azimuth encoded in our model (i.e., the number of elevations in the ‘OTid’ space map).
- *N* = number of model Imc-like neurons, i.e., neurons with Imc-like anatomical projection patterns.
- *k_max_* = maximum number of RF lobes allowed for each model neuron.
- *N\** = smallest number of upto-k_max_-lobed model neurons needed to solve location-invariant selection across *L* elevations.

The optimization model solves for the number and positions of RF lobes of each of the *N* model neurons in order to achieve location-invariant selection. The model neurons are ‘Imc-like’: each of them is excited by a stimulus placed anywhere within its RF, and delivers competitive inhibition to all locations in the OTid space map outside its RF that is proportional to the strength of the stimulus (Supplementary Fig. 4ab). Without loss of generality, we take stimulus priority = 1 unit (for all stimuli), and the proportionality constant (underlying inhibition by the Imc) to be 1. Therefore, for each stimulus, each neuron excited by that stimulus generates an inhibition of 1 unit at those locations in the OTid that are outside that neuron’s RF (Supplementary Fig. 4ab). For successful, relative-priority dependent competitive stimulus selection between stimuli presented at a given pair of locations, the net inhibition at these two locations in the OTid should be equal. For location-invariant competitive selection, this condition must hold for stimuli placed at any pair of all the possible *^L^c_2_* (L choose 2) pairs of locations. The details of the setup of the optimization problem are described below.

Let *X* be a matrix of size *L X N* (Supplementary Fig. 4c), where the *j^th^* column of the matrix corresponds to the *L* elevational locations encoded by the *j^th^* Imc neuron in the population. The optimization problem is framed as *min_X_* f (*X; L, N*), where the objective function f(*X*) is designed such that it achieves its minimum value (of *-L(L-1)*) for a given *L* only when the RFs of the model neurons achieve location-invariant selection.

Consider two competing stimuli (of equal strength) placed at locations 1 and 2. In our scheme, we represent this by a row vector ***a***_1xn_= [1 1 0 ….0 …0] (Supplementary Fig. 4d). The ones in the first two indices of the row vector correspond to the two locations at which the competing stimuli are placed.

Note that *X^T^**a**^T^* results in a vector in which the *j^th^* index corresponds to the number of locations that the *j^th^* neuron is activated by when the two competing stimuli are placed in positions shown in **a** (Supplementary Fig. 4e).

Additionally, the matrix (X-1) corresponds to the suppression image of the Imc population. The *j^th^* column of this matrix represents the locations to which the *j^th^* Imc neuron sends inhibition in the OT space map. This is because of the inverse anatomical projections from the Imc to the OT. The product *(X-1)X^T^**a**^T^* then results in a vector in which the *j^th^* index corresponds to the net inhibition sent to the *j^th^* location by the entire Imc population when the two competing stimuli are placed at different locations, i.e., at different positions within the row vector ***a*** (Supplementary Fig. 4f).

For competitive selection at these two locations, the net inhibition at these two locations in the space map of the model ‘OTid’ should be equal. To penalize solutions for which this is not the case, we include a cost term in the objective function that is equal to the square of difference in the inhibition at the two locations. This is written mathematically as

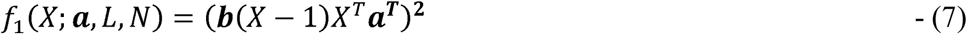

where ***b*** is a row vector whose length equals that of ***a*** and nonzero indices are same as a, but with the sign of one of the 1s flipped (in this case ***b***= [1 −1 0 ….0 …0] or [-1 1 0 ….0 …0]). The minimum value that *f_1_* can take is 0, which happens when equal inhibition is sent to both the locations at which the competing stimuli are placed (Supplementary Fig. 4f).

In addition to the strength of inhibition at the two locations being equal, the strength of inhibition must be strictly negative. This is because, the other possibility, of strength of inhibition at each location being zero, would not be acceptable because no inhibition would be sent to either of the two locations. To penalize solutions for which this condition is not met, we include a cost term in the objective function that is equal to the number of locations at which the inhibition is not negative. This is written mathematically as

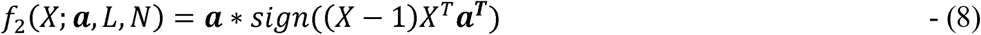

Minimizing *f_2_*, therefore, ensures that inhibition is sent to both the locations. The minimum value *f*_2_ can take is −2, when inhibition is sent to both the competing locations (Supplementary Fig. 4f).

Finally, we write the full objective for the location-pair (specified via vector **a**) as

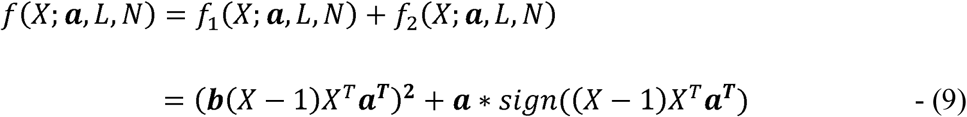

the minimum possible value of which is −2.

For location invariance to be achieved, the function *f* should be minimized for each pair of locations at which competing stimuli can be placed. In other words, *f* should be minimized for all possible permutations of vector **a**. This can be written mathematically as

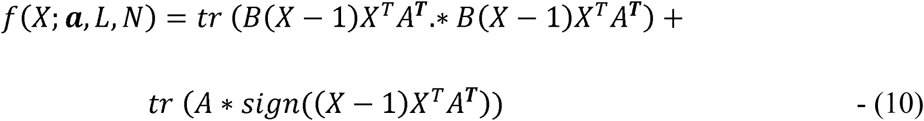

where *A* is the permutation matrix of **a** for all possible location-pairs and *B* is the corresponding permutation matrix of **b**. *tr(Y)* refers to the trace (sum of all the diagonal elements) of the matrix *Y*, Y.*\*Z* is the Hadamard (element-wise) product between the matrices *Y* and *Z* and *sign(Y*) is a matrix obtained by applying the element-wise sign operator to the matrix *Y*.

Because there are *^L^c_2_* possible location-pairs (corresponding to the *^L^c_2_* permutations of the vector **a**), the minimum value that *f* can achieve is −2\**^L^c_2_ = −L(L-1)*. Thus, location-invariant selection is achieved in our optimization model if and only if the cost function converges to the lowest possible value of *–L(L-1)*.

We add two constraints to this optimization scheme. First, we code the RFs of all the model neurons with ones (inside RF) and zeros (outside RF), a simplifying assumption (see “Model assumptions” section below for implications of this assumption). Second, we introduce a mechanism to limit the number of lobes in any model neuron to *k_max_*. This is done so that, by setting *k_max_* = 3, we would be able to match the experimentally observed constraint that there are no more than three RF lobes per Imc neuron. The first constraint is fed into the optimization problem as bounded integer constraints with bounds between 0 and 1 to make the RFs binary. The second constraint is implemented as an inequality constraint, written mathematically as

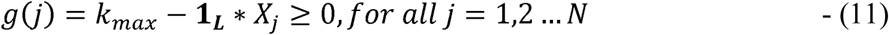

where **1_L_** is a row vector of length *L*, and *X_j_* is the *j^th^* column of *X* corresponding to the RF of the *j^th^* neuron. Additionally, we also test the model with *k_max_* = 10 for some of the analyses reported in Fig. 4, Fig. 6 and Supplementary Fig. 5.

We solve the above nonlinear optimization problem with mixed constraints, an np-complete problem, using the *‘MIDACO’* solver in MATLAB ^37^.

#### Estimating *N\**

*N\** is the smallest number of model neurons needed to solve location-invariant selection for a given *L* and *k_max_*, i.e., the smallest *N* for which the minimum value of the objective function (*-L(L-1)*) can be successfully achieved. This was estimated as follows. For each value of *N* from 1 to *L*, we ran the optimization model 1000 times (1000 runs). Any given run was said to have converged to a solution if the value of cost function did not change for 1000 successive iterations (by setting the *‘evalstop’* criterion in the optimization code to 1000), thereby reaching an asymptotic value. The collection of model neuron RFs at convergence was called a ‘convergent solution’. Additionally, if the convergent solution attained the value of *–L(L-1)*, then it was called an ‘optimal solution’. In other words, optimal solutions are ones that converged and additionally achieve location-invariant stimulus selection.

*N*\* (for a given *L* and *k_max_*) is, therefore, the smallest value of *N* for which at least one of the 1000 runs yielded an optimal solution, meaning that for *N = N\**-1, none of the 1000 runs yielded a solution that successfully achieved location-invariant selection.

For instance, if *k_max_=*1 lobe, then for all *L*, *N* = L* (Fig. 4d, blue data; consistent with theoretical calculation presented in the text surrounding Fig. 3). If *k_max_* = 3 lobes and *L* = 5 elevations, all runs for all values of N from 1 to *L* yielded convergent solutions, but *optimal* solutions were produced only when *N* ≥ 4 (Supplementary Fig. 5a). More generally, if *k_max_* > 1 lobe, then for all *L* > 4, *N* < L* (Fig. 4d; red and black data).

#### Range of *k_max_* values chosen for various analyses (Fig. 4d onwards)

The specific values of *k_max_* used in our simulations (Fig. 4 and 6) were 1, 3, and 10 lobes. The reasoning for this choice of values is described below.

- *k_max_* = 1 lobe corresponded to the null hypothesis of single lobed RFs
- *k_max_* = 3 lobes represented Imc data (Fig. 1i)
- *k_max_ =10* lobes. (i) The range of elevations encoded by the OTid and the Imc is no greater than −60° to −60°, and (ii) Most individual RF-lobes have a half-max height ≥ 10° (10-percentile value of half-max height of an individual RF lobe = 10° (Supplementary Fig. 3e). Therefore, the number of possible distinct lobes along elevation for RFs of typical Imc neurons ≤ ∼10 lobes (=120°/ (10° + 2°); with the two added degrees representing 1° spacing on either side of a lobe to separate it from abutting ones.)

#### Model assumptions

Our optimization model makes two key simplifying assumptions: (a) discretized (pixelated) spatial locations, and (b) binary (on or off) RFs of the model neurons. The former assumption can be readily reconciled with biology by making the pixel size sufficiently small. Therefore, this assumption does not result in loss of generality of the model. Second, the pattern of spatial inhibition sent to the OTid space map, the key computational function required of Imc in the model, is the spatial inverse of the RF: inhibition is sent to all locations except the ones inside the RF. In other words, the spatial pattern of inhibition is, by definition, a ‘*binarized* spatial inverse’ of the Imc RF, with the strength of delivered inhibition being proportional to the specific location within the continuous RF at which the stimulus is placed (Supplementary Fig. 4ab). For the model, it is the pattern of inhibition that is critical, informationally speaking, rather than the variations in the strength of delivered inhibition based on the specific location within RF that a stimulus occupies (Supplementary Fig. 4ab). (This is unlike population vector coding, where the specific values of firing rates within an RF are critical informationally ^25–28^). Therefore, the continuous RF can be binarized itself (say, at the half-max, or 75%-max level) without the qualitative conclusions of the model being affected (Supplementary Fig. 4ab). Notably, despite these simplifying abstractions of the biology by the model, we found that predictions from the model held true experimentally (Fig. 5), further revealing that the model captured sufficiently well the key computational principles at play in this circuit. Consequently, it was able to provide a compelling explanation for the unusual functional properties of Imc neurons, and illuminate neural mechanisms of location-invariant stimulus selection in this midbrain circuit.

### Characterizing signature properties of optimal model solutions, and testing them in experimental data (Figs. 4 and 5)

#### The “multilobe property” (property #1)

<u>Model:</u> For each optimal solution at each (L,N*,k_max_) tested, we examined if any of the model RFs were multilobed. A model RF was said to be multilobed if it had “on” pixels that were separated by “off” pixels; two adjacent “on” pixels were treated as one lobe. For instance, in Fig. 4a, neurons #2 and #4 have one lobe each. Neuron #1 has two RF lobes and neuron #2 has 3 RF lobes. These two neurons are multilobed. Thus, this optimal model solution is said to satisfy the “multilobe property”.

<u>Data:</u> For each coronal Imc plane recorded, we examined if any of the neurons in that plane had multilobed RFs. Whether an RF was single or multilobed was determined using methods described in (and surrounding) Fig. 1.

#### The “optimized lobe-overlap property” (property #2)

<u>Model</u>: A multilobed model neuron that shares each of its RF lobes, but not all, with another neuron is said to satisfy this property. If every neuron in a model solution satisfies this property, the model solution itself is said to satisfy the optimized lobe-overlap property. The fraction of model solutions satisfying this property for each (L, N*) is plotted in Supplementary Fig. 5C (100%, in each case).

<u>Data</u>: The set of neurons recorded within a given coronal plane, i.e., across all the recording sites along a dorsoventral penetration, is collectively a potential solution set for location-invariant selection across all elevation pairs at that azimuth. (This is because of our finding that spatial azimuth is encoded topographically along the rostrocaudal axis of the Imc, and all the elevations at a given azimuth are encoded by the neurons in the coronal plane at the appropriate point along the rostrocaudal axis; Fig. 2 and Supplementary Fig. 3). A multilobe neuron that shares at least one of its RF lobes, but not all, with another neuron in the solution set is said to satisfy the experimentally testable version of the lobe-overlap property. To test this property in data, we first obtained the set of discrete elevational locations encoded by Imc neurons in a solution set (coronal plane). We did this by quantizing, at a resolution of 3° (to match theory and model; see main text related to Fig. 3), the maximum elevation range encoded by their RFs combined. Next, an RF lobe of a multilobed Imc neuron was said to overlap with the RF of another neuron if there existed a location within the former’s half-max extent that also lay within the half-max extent of the latter’s RF. The fraction of multilobed Imc RFs in each coronal plane that satisfy this testable version of the optimized lobe-overlap property is shown in Fig. 5c. (This testable version of the lobe-overlap property was necessary because of the inherent infeasibility of recording from all Imc neurons in a coronal section, i.e., from all the neurons in a ‘solution set’. Specifically, the small ML extent of the Imc (<350 µm), coupled with the thickness of the electrode (250 µm) that was used to reliably target the deep Imc (∼16 mm below brain surface), limited us to one dorsoventral penetration within a coronal section. This made recording from all Imc neurons in a given section unviable. The average # neurons recorded per section = 3.44 ± 0.47.

#### The ‘combinatorial’ property <u>(property #3).</u>

<u>(A) *“Assorted neural subset” feature*</u>: Distant neurons are recruited to achieve selection for nearby locations, and nearby neurons are recruited to achieve selection for distant locations. To test for this feature, we divide the elevation range (L locations) into three parts, the upper L/3, middle L/3 and lower L/3 locations. Two locations are said to be ‘*nearby’* if the distance between them is ≤ L/3, and ‘*distant’* if the distance between them is ≥ 2*L/3. Similarly, two neurons are said to be nearby if the distance between them is ≤ (N-1)/3, and distant, if their distance is ≥ 2*(N-1)/3. We then ask if distant neurons are recruited for a nearby location-pair (LP), and vice-versa. Since there is no meaningful functional ordering of multilobe neurons owing to the lack of topography in the encoding of elevation, we must test these questions across permutations of the ordering of Imc neurons within a solution.

<u>Model</u>: First, we tested if distant neurons are recruited for a nearby location-pair. We did so by computing the following metric (eq. (12)) for each (L, N*) (Fig. 4f).

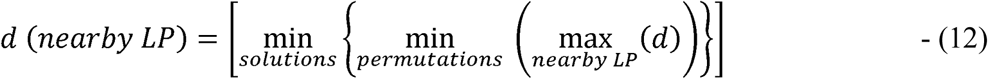

Here, *‘d’* is the maximum distance between the neurons recruited for solving selection for a given nearby location-pair in a given solution. The maximum of this across all nearby location-pairs yields the farthest distance between neurons recruited to solve selection for any nearby location-pair. The minimum of this value across permutations of neurons in the solution, and across all solutions, yields *d (nearby LP)* for that (L, N*).

For L=5 (N*=4), we tested this exhaustively for all possible permutations (*4!*). However, for L = 20 (N* = 14) and L = 40 (N* = 27), the number of permutations is very large (*14!* = 8.7 x10^10^ and *27!* = 1.08 x 10^28^). Because it was infeasible to test all possible permutations in these cases, we tested a subset of permutations (n=1000) that was selected randomly from the set of all the possible permutations using the *‘randperm’* function in MATLAB.

For each (L, N*), we calculated the normalized minimum distance between neurons recruited for selection at distant location-pairs as shown in eq. (13), and plotted it in Supplementary Fig. 5e.

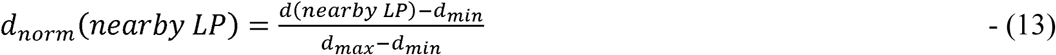

Here, *d_max_* (= N*-1) and *d_min_* (= 1) are the maximum and minimum possible distances between neurons in a solutions set consisting of N* neurons. We found that in every case, this normalized distance was high (>0.66; the normalized cut-off value chosen for defining ‘distant’ neurons).

Next, we tested if nearby neurons are recruited for a distant location-pair, using a metric constructed with a logic similar to that used above:

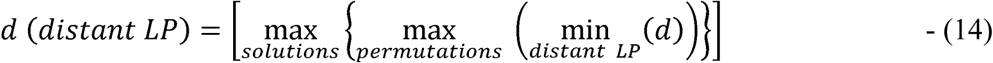

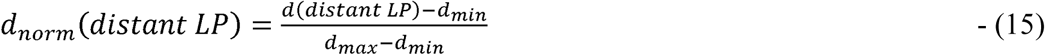

For each (L, N*), we calculated the normalized maximum distance between neurons recruited for selection at distant LPs (eq. (15)), and plotted the results in Supplementary Fig. 5f. We found that in every case, this normalized distance was small (<0.33; the normalized cut-off value chosen for defining ‘nearby’ neurons).

<u>Data</u>: For Imc neurons in each solution set (coronal plane), we obtained the range of discretized elevation values encoded as before (resolution of 3°), and then calculated the normalized minimum distance between nearby neurons and the normalized maximum distance between distant neurons using the Eq. (13) and (15) above. Note that for the notions of nearby neurons and distant neurons, there need to be at least 3 neurons in the solution set so that the maximum distance is 2 and the minimum distance is 1. Out of 14 coronal planes that contained multilobe 3 neurons. The results from these 8 planes are plotted in Fig. 5f.

<u>(B) *“Extensive intersection” feature*</u>. Location-pairs occupying distant portions of space recruit shared neurons to solve selection at each pair. Two location-pairs are said occupy distant portions of (elevational) space if one location-pair lies within the upper third of the locations (upper L/3) and the other lies within the lower third of the locations (lower L/3). Since intersection between the neural subsets is independent of the ordering of the neurons, we do not need to test this for all permutations of neuron orderings. <u>Model</u>: For every optimal solution at a given (L, N*), we tested if there existed two location-pairs in distant portions of space such that the neural subsets recruited to solve selection at each location-pair shared at least one neuron. The fraction of optimal solutions that satisfied this property is plotted as a function of (L, N*) in Supplementary Fig. 5g; the fraction is uniformly 100%.

<u>Data</u>: For Imc neurons in each solution set (coronal plane), we obtained the range of discretized elevation values encoded as before (resolution of 3°). We then tested if these neurons satisfied the extensive-intersection property as described for the model. Of the 14 coronal planes at which neurons were recorded, in 6 cases, the encoded locations included two location-pairs that occupied distant locations. The fraction of these 6 coronal planes that satisfied the extensive intersection property is shown in Fig. 5g (100%).

### Wiring and metabolic costs of location-invariant selection in the Imc-OT circuit (Fig. 6)

<u>Wiring cost</u>: The wiring cost for location-invariant selection by the Imc is estimated as the cost of generating axonal projections (‘wires’) between each Imc neuron and each of its target OTid neurons. This cost depends both on the number of locations that each neuron must suppress and the number of neurons in the population. Assuming that the lengths of wires from Imc to each OTid neuron is approximately equal (say 1 unit each, without loss of generality), we can estimate the total wiring length and consequently the total wiring cost using Eq. (16) below (see ^22^).

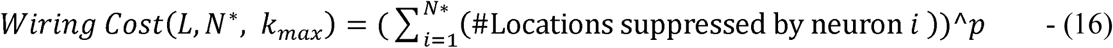

The summation is the total wiring length of all the wires from the Imc neurons to the OTid population. *‘p’* is a power term such that typically 1 < *p* < 4 (see ^22^). This quantity is computed for each optimal solution (obtained over the 1000 runs) for a given (L, N*, k_max_) triplet, and the results are plotted in Fig. 6a.

<u>Metabolic cost</u>: The metabolic cost for location-invariant selection by the Imc is estimated as the cost of generating and broadcasting spikes to the OTid to achieve competitive suppression. This depends on the number of neurons activated by a stimulus at each of the L locations, as well as the number of OTid locations to which each activated neuron delivers inhibition. If the cost of suppressing one OTid location using 1 spike is 1 unit, then the total metabolic cost for the circuit for a given firing rate *f* is given by Eq. (17) below (using a similar formula as for wiring cost).

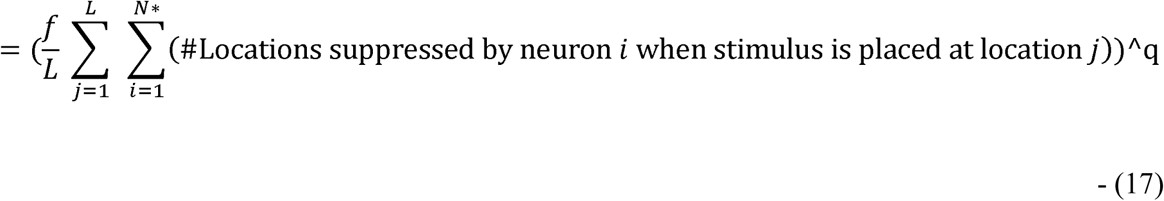

Note that the term in the inner summation is non-zero only for activated neurons when the stimulus is placed at location *j*. *‘q’* is a power term chosen such that 1 < q <4 (similar to the wiring cost). This quantity is computed for each optimal solution (obtained over the 1000 runs) for a given (*L, N*, k_max,_ f = 10 Hz*), and the results plotted in Fig. 6b.

<u>Total cost</u>: The total cost for any solution is calculated as a weighted combination of the wiring cost (weight = *α*) and the metabolic cost (weight = *β*) as given in Eq. (18) below. There are five parameters in this summation (*α, p, β, q* and *f*). The results are plotted for *α = 20, p = 2.5, β = 80, q= 2.42* for firing rates of *f*=10 Hz (blue curve) and *f*=80 Hz (red curve) in Fig. 6c.

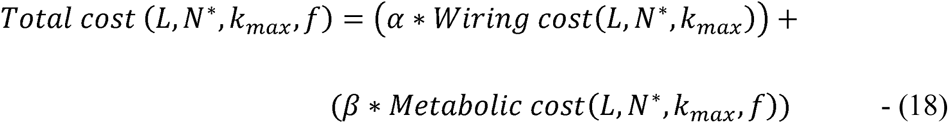

### Data analyses and statistical tests

All analyses were carried out with custom MATLAB code. Parametric or non-parametric statistical tests were applied based on whether the distributions being compared were Gaussian or not, respectively (Lilliefors test of normality). The Holm-Bonferroni correction was used to account for multiple comparisons. Data shown as a ± b refer to mean ± s.e.m, unless specified otherwise. The ‘*’ symbol indicates significance at the 0.05 level (after corrections for multiple comparisons, if applicable). Correlations between RF centers (azimuth) and electrode measurement positions (rostrocaudal/ dorsoventral) were tested using Spearman’s rank correlation coefficient (*corr* command in MATLAB with the Spearman option).

### Code and data availability

Software code and the data that support the findings of this study are available from the corresponding author upon reasonable request.

## Supplementary Figures

**Supplementary Fig. 1.**
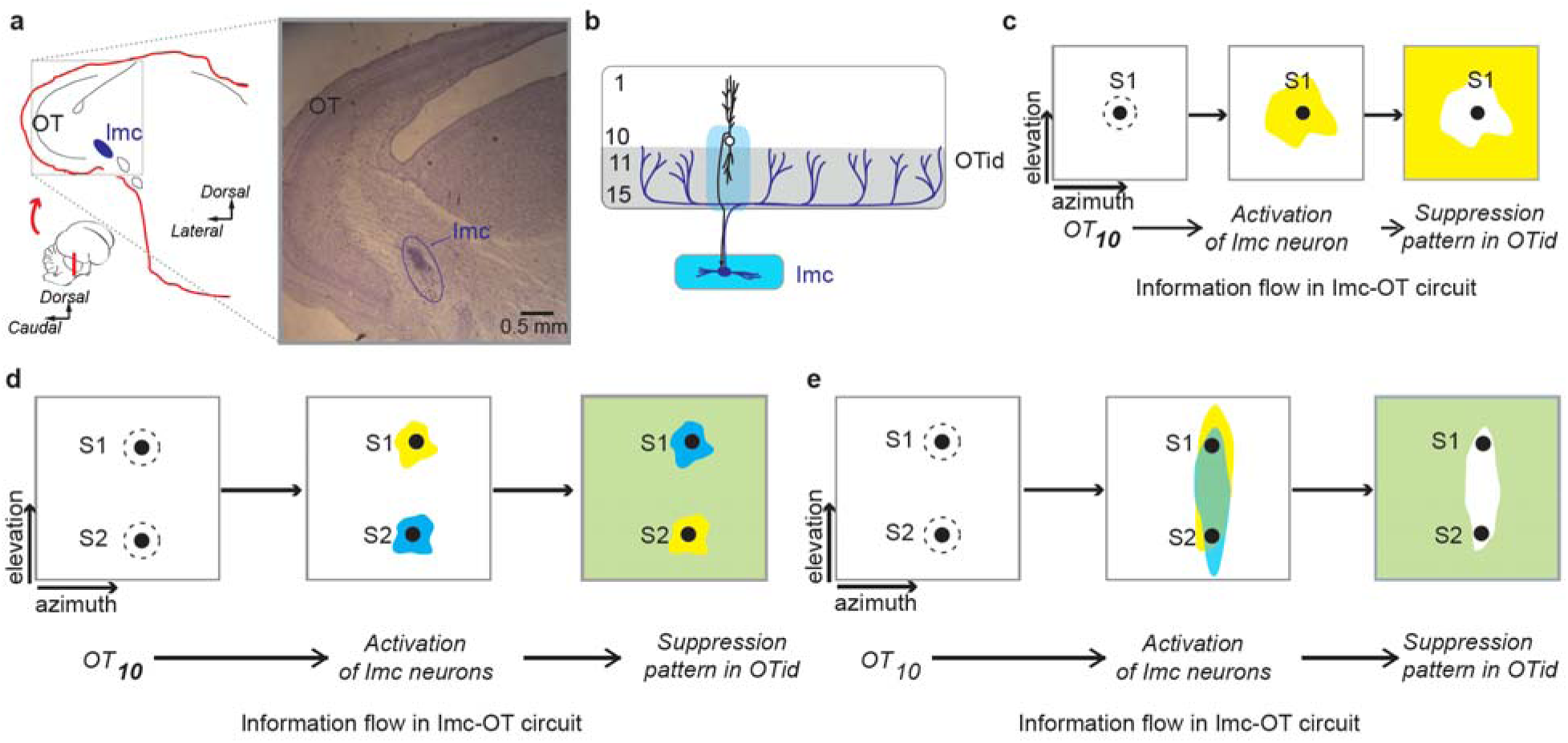
Anatomical connectivity and information flow between the Imc and optic tectum (OT). Related to Fig. 1. **(a)** *Left:* Cartoon showing side view of barn owl brain (inset), and coronal section taken along the indicated line in inset. *Right*: Nissl stained, coronal section of midbrain depicting the multilayered optic tectum (OT) and the isthmi pars magnocellularis (Imc). The OT_10_ is seen as a darkly stained arc of cell bodies. The Imc is a long m long rostrocaudally and 350 appears in transverse sections as a 700-μm x 350-μm elliptical disk of neurons (blue oval) ^16^. The long, rostrocaudal axis of the Imc is parallel to the rostrocaudal axis of the OT. Dark area in the dorsal portion of Imc: electrolytic lesion following stereotactic and electrophysiologically-based targeting of Imc. **(b)** Schematic of anatomical connectivity between the Imc and OT. Imc neurons receive input from a focal portion of OT_10_ (black neuron), but project broadly back (blue lines) to the OTid sparing just the portion of the space map providing input (light blue shading across OT layers) ^6^. All layers of OT are known to represent space topographically, but how the Imc represents space is not well understood (see also (e)). **(c)** Schematic of information flow through the OT_10_-Imc-OTid circuit showing the functional, spatial-inverse operation executed by established Imc-OT connectivity ^6^. Maps of visual space in the OT_10_ (*left*), Imc (*middle*) and OTid (*right*). For purposes of illustrating the spatial inverse operation, Imc RFs are assumed to be large with an unknown shape (yellow shading). A visual stimulus S1 at location 1 activates the space map in OT_10_ (left; dashed circle - RF of activated neuron). This, in turn, activates an Imc neuron (middle; yellow represents assumed RF of activated Imc neuron), which delivers inhibition to all locations in the OTid space map that are outside the RF of the activated Imc neuron (right; yellow shading) ^6, 15, 16^. **(d)** Schematic representation of stimulus selection in the OTid under the assumption that Imc RFs are small, resembling OT_10_ RFs. *Left*: Shown are two stimuli S1 and S2, at locations 1 and 2, respectively, which activate corresponding neurons in the OT_10_ space map. *Middle*: Imc neuron activated by S1 (yellow RF), and Imc neuron activated by S2 (blue RF). *Right*: Combined pattern of suppression generated in the OTid by the activated Imc neurons: each neuron delivers suppression to locations outside its RF; green = yellow + blue. Each stimulus successfully suppresses the other – S2 lies within the yellow zone of suppression produced by S1, and vice-versa – implementing selection for stimuli at these two locations. Similarly, if every spatial location was encoded by an Imc neuron with a small, OT_10_ like RF (and with space-inverting connectivity with the OT), then stimulus selection in the OTid would be achieved successfully for all pairs of locations (the ‘copy-and-paste’ strategy described in the text). (**e**) Same as (d), but with Imc RFs that are large and elongated vertically, covering almost the entire elevational extent, as reported in the literature ^17, 18^. Shown in the middle panel are the RFs of two Imc neurons, in yellow and blue, respectively. *Left*: As in (d). *Middle*: S1 activates both Imc neurons, and so does S2. *Right*: Resulting patterns of inhibition in the OTid space map; green = yellow + blue; large swaths of space are left without inhibition (white region in right panel, corresponds to intersection of the two RFs). Specifically, neither stimulus is suppressed by the other even though the two stimuli are well separated in elevation (shown here to be approximately 60° apart), preventing stimulus selection along the elevation. Large, vertically elongated Imc RFs, therefore, are unable to support spatial selection in the OTid across all elevational locations. This is an apparent paradox in terms of Imc-OT function because the OTid is known to exhibit location-invariant selection, including when stimuli are <10° apart in azimuth or elevation ^10, 12–14^, with Imc driving this global competitive selection ^16^.

**Supplementary Fig. 2.**
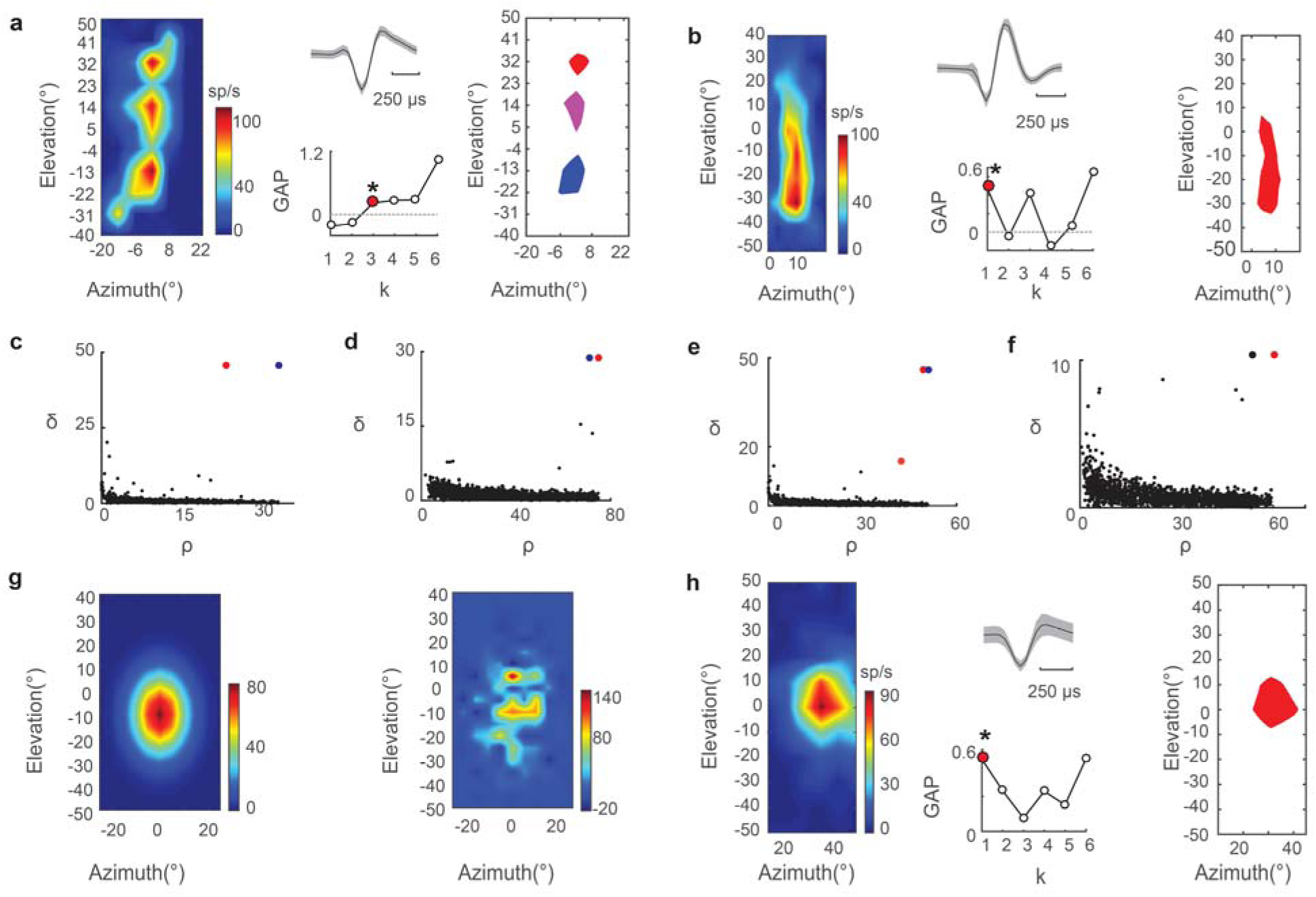
Analysis of visual RFs of example Imc and OTid neurons. Related to Fig. 1. **(a)** Three-lobed visual RF of an example Imc neuron. (*Left*) Color coded rate map of RF. (*Middle, top*) spike waveform for the neuron. (*Middle, bottom*) GAP statistic plot. (*Right*) Half-max extents of the 3 lobes identified by model selection with the gap statistic. **(b)** Single-lobed visual RF of an Imc neuron; conventions same as (a). **(c)** Density peaks clustering method. Scatter plot of local density (ρ) around each datapoint in Fig. 1c vs. the distance of that datapoint from other points that have higher local density (δ). (For the point with highest local density, δ is conventionally taken as the maximum distance of the point from all other points). Points that have both high local density (large ρ value) and that are far away from other points of high local density (large δ value) are potential cluster centers; Red and blue points in this example. Red point corresponds to the center of top cluster, and blue point, the center of lower cluster shown in Fig. 1d. **(d-f)** Same as (c), but for RFs in Figs. 1f, Supplementary Figs. 2a and 2b respectively. **(g)** Effect of sampling resolution and neuronal noise on detection of optimal number of lobes in the data. (*Left*) The simulated single-lobed 2D Gaussian RF used for the Monte-Carlo analysis in Fig. 1j (Methods). Shown are mean firing rates at different locations. (*Right*) Plot of the RF obtained when it is re-simulated after adding noise (Fano-factor = 30), and sampled with step-sizes = 5° in azimuth and elevation. This sampled RF was identified as having two lobes by our analysis pipeline (conversion to distribution of points on plane, density peak clustering, followed by gap statistic model selection), which is incorrect because the true underlying Gaussian RF was single-lobed. This illustrates how noisy neural responses can lead to the erroneous conclusion that a single-lobed RF is multilobed. **(h)** 2D visual RF of an example OTid neuron. Conventions as in (a), (b). The RF is single lobed.

**Supplementary Fig. 3.**
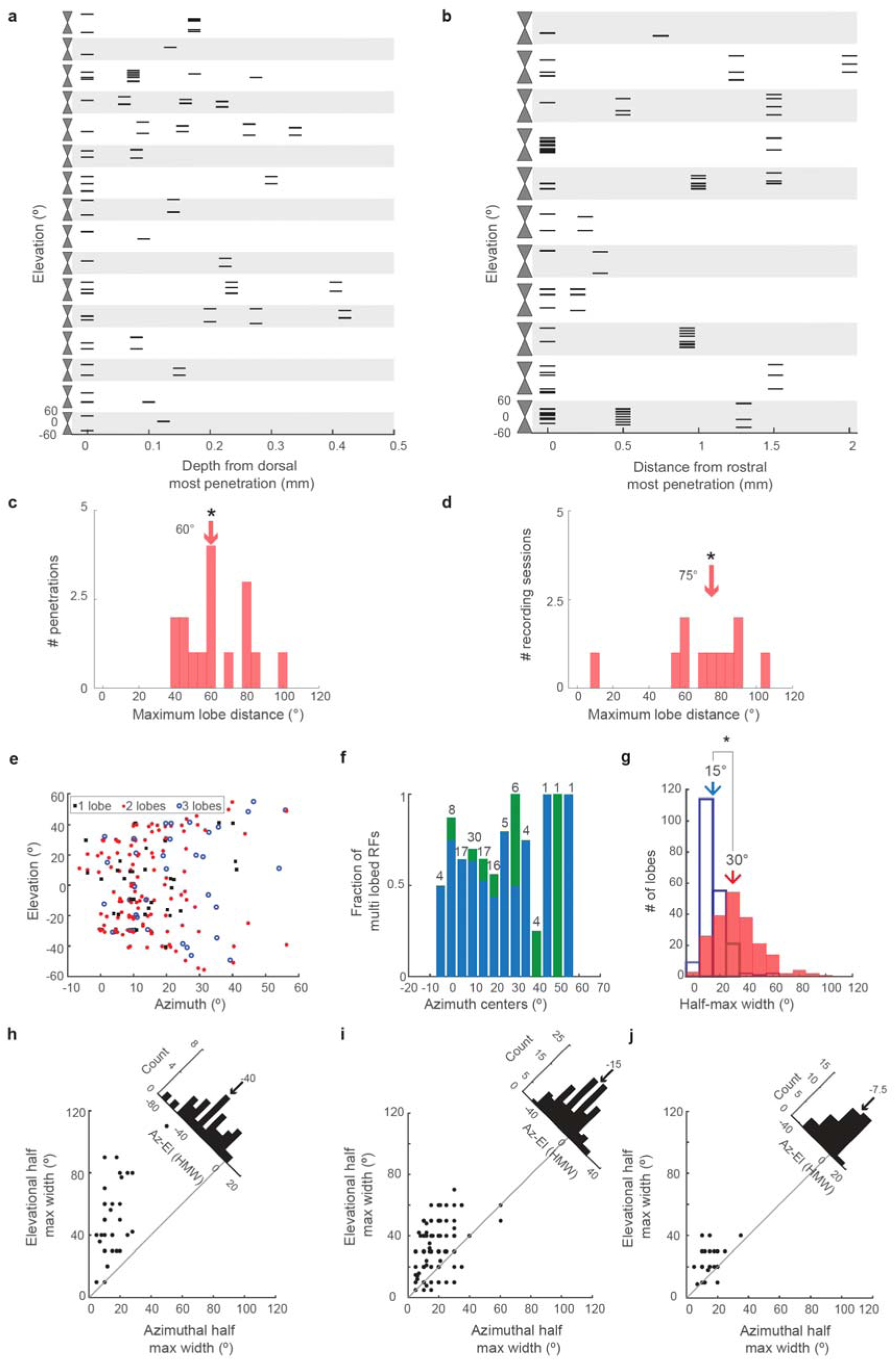
Detailed analysis of the organization and structure of RF lobes of Imc neurons. . Related to Fig. 2. **(a)** Plot of elevational centers (black horizontal ticks) of all the visual RF lobes of all individual neurons recorded at a multiunit site, as a function of the dorsoventral position of the electrode within the Imc along a penetration. Each horizontal band (gray and white) band depicts a different penetration; the vertical extent of each band spans −60° to +60° in elevation. No systematic organization of elevational centers of RF lobes along the dorsoventral as evidenced by widespread and irregular distribution of lobe centers at each depth within a penetration. **(b)** Plot of elevational centers (black horizontal ticks) of all the visual RF lobes of all individual neurons recorded at a multiunit site, as a function of the rostrocaudal position of the electrode during that recording session. Each horizontal band (gray and white) depicts a different recording session; the vertical extent of each band spans −60° to +60° in elevation. No systematic organization of elevational centers of RF lobes along the rostrocaudal axis, as evidenced by widespread and irregular distribution of lobe centers at each penetration (within a recording session). (**c**) Histogram showing maximum distance between RF lobes measured along each penetration (i.e., each horizontal band in (a)). * indicates mean significantly different from 0 (p < 0.001); one tailed t-test. (**d**) Histogram showing maximum distance between RF lobes measured across all penetrations made along the rostrocaudal axis in each recording session (i.e., each horizontal band in (b)). * indicates mean significantly different from 0 (p < 0.001); one tailed t-test. **(e, f)** Multilobe neurons were found at all tested azimuths. (e) Scatter plot of the azimuthal and elevation centers of the individual lobes of multilobed RFs of all neurons. (f) Fraction of measured RFs that were multilobed, plotted as a function of the azimuthal center of the RF (blue corresponds to 2-lobed Imc RFs and green to 3-lobed Imc RF). (**g**) RF lobes of Imc neurons are elongated in elevation. Histogram of azimuthal (open) and elevational (red) half-max widths of all the RF lobes across all recorded neurons. Arrows indicate median values. * indicated that lobes are larger in elevation than in azimuth (ranksum test p < 10^-25^). (**h, i, j**) Lobes of single-lobed RFs are taller (more elongated in elevation) than those of two-lobed RFs, which are in turn taller than those of three-lobed RFs. Scatter plot of elevational vs. azimuthal half-max widths of single-lobed RFs (h), two-lobed RFs (i), and three-lobed RFs (j). *<u>Insets</u>*: Histogram of data points projected onto the line perpendicular to the line of unity. The median values of the histograms increase (and approach zero) as we go from panel h (single-lobed RFs) to j (three-lobed RFs). HMW: Half-max width.

**Supplementary Fig. 4.**
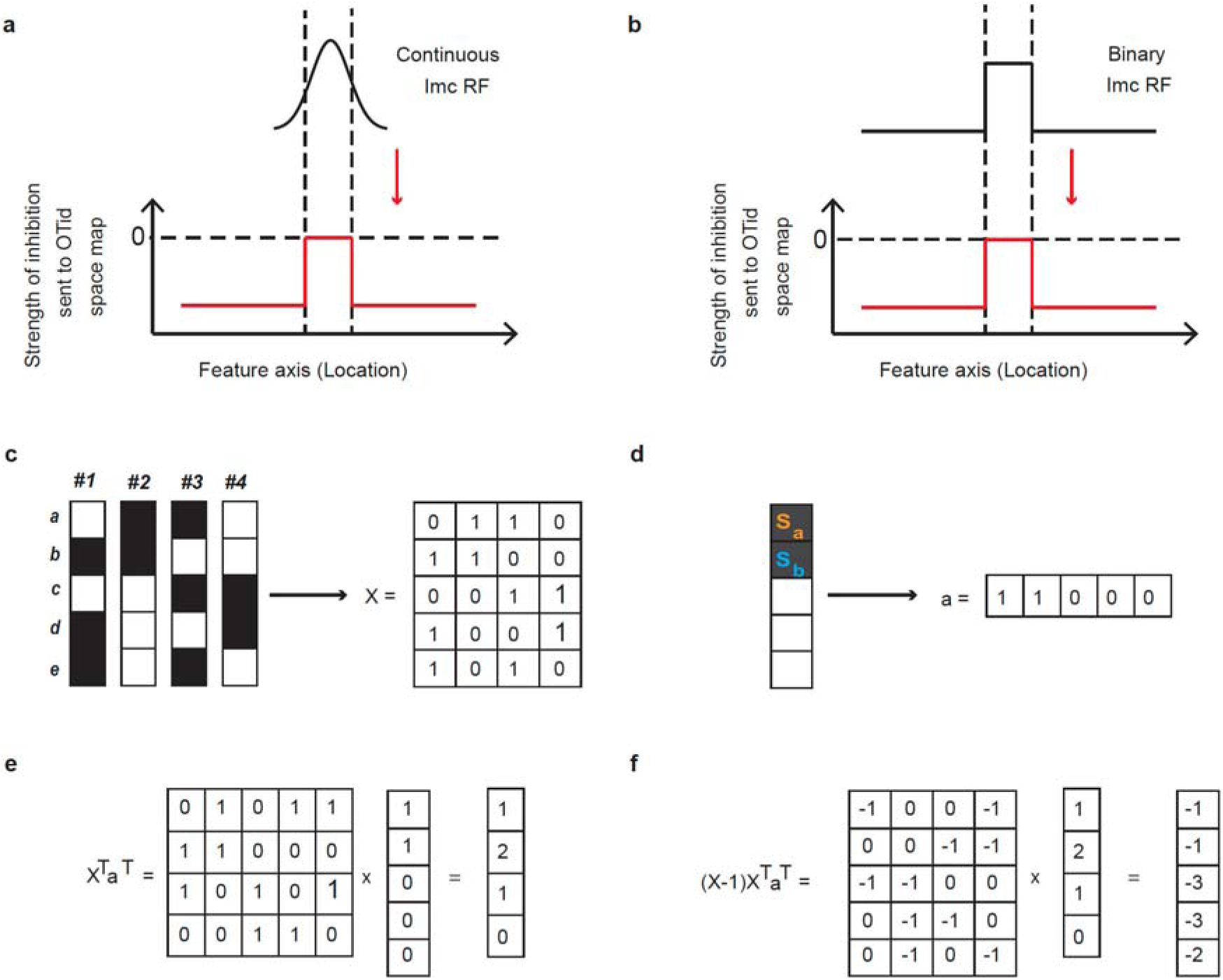
Setup of location-invariant stimulus selection as an optimization problem. Related to Fig. 4. **(a-b)** Patterns of spatial inhibition sent by the Imc to OT by continuous Imc RFs (biologically-consistent), vs. binary Imc RFs (simplified abstraction for modeling purposes). (a) *<u>Top</u>:* Schematic of an Imc RF that encodes locations using continuous values of firing rates. *<u>Bottom:</u>* Pattern of inhibition sent by the Imc RF to the OTid space map based on the space inverting anatomy between the Imc and OT. Without loss of generality, locations outside the half-max extent of the Imc RF are considered to be spared by Imc spatial inhibition. (b) Same as (a), but when the Imc RF is assumed to be binary at the half-max level of the RF (Methods). The spatial pattern of Imc inhibition in (b) is nearly identical to that in (a) (with the exception that the strength of inhibition in a gets scaled based on the specific position of the stimulus within the RF half-max.) **(c)** *<u>Left</u>*: RF solutions (from Fig. 4a) obtained by the optimization problem when L=5 and N=4. *<u>Right</u>*: Same RFs represented as an L *x* N matrix *‘X’* for the optimization problem. **(d)** *<u>Left</u>*: Stimuli presented at locations *a* and *b* (from Fig. 4b). *<u>Right</u>*: Stimuli pair represented as a row vector for the optimization problem. **(e)** The product X^T^a^T^ results in a vector of length N *X* 1 whose *j^th^* element equals the number of locations that activate model neuron *j.* For instance, neuron #2 is activated by both S_a_ and S_b_. So, the second element of the vector X^T^a^T^ is 2. **(f)** The product (X-1)X^T^a^T^ results in a vector of length Lx1 whose *j^th^* element equals the net inhibition sent by the Imc population to location *j* when the stimuli are presented at locations indicated by vector a. For instance, the inhibition sent to location b is −1 (by Imc neuron #3). So the second element of (X-1)X^T^a^T^ is −1.

**Supplementary Fig. 5.**
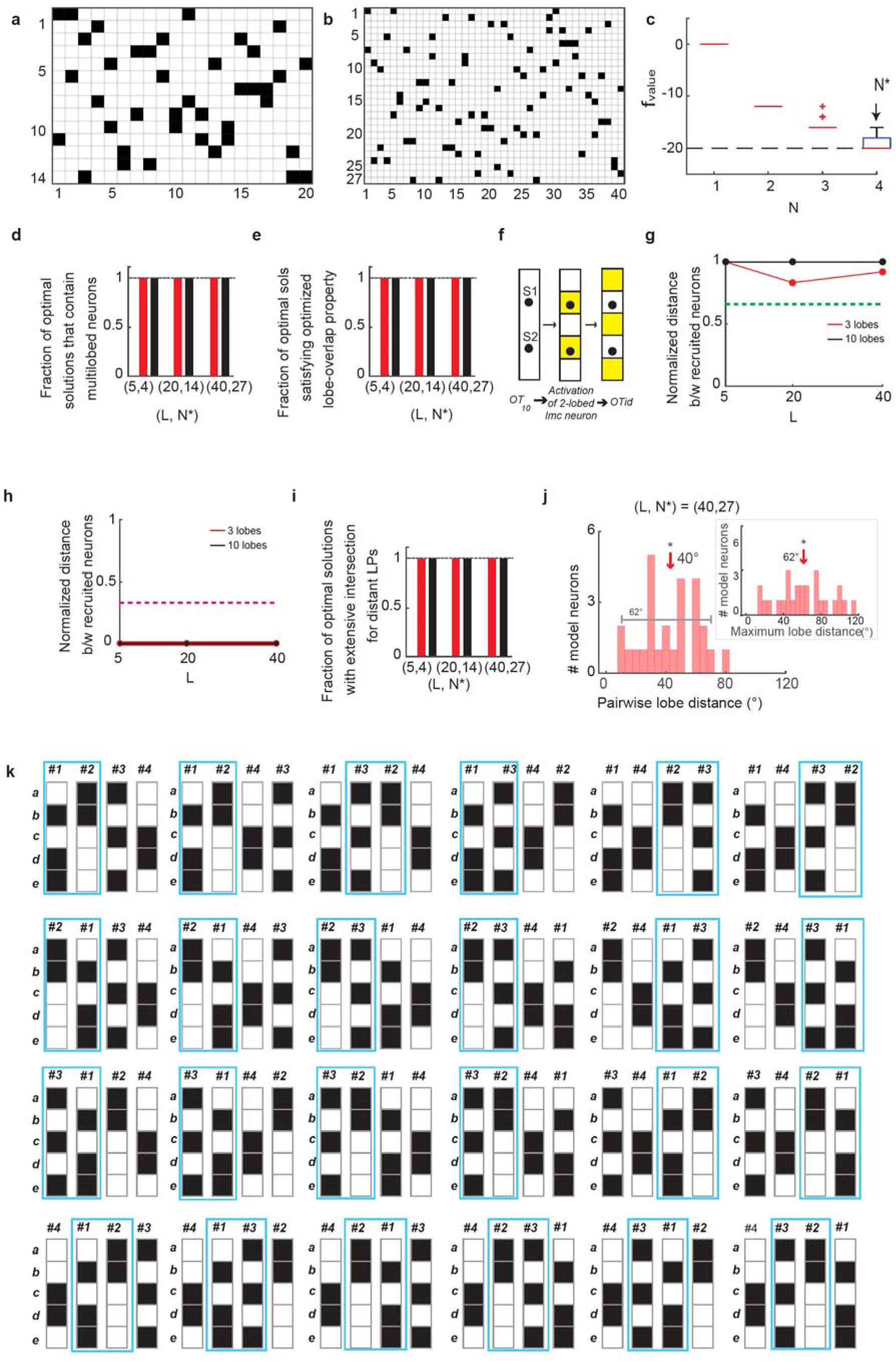
All optimal model solutions exhibit signature properties of combinatorially optimized inhibition. Related to Fig. 4. **(a)** Example optimal solution for (L, N*) = (20, 14). Black pixels: Locations inside neurons’ RF, White pixels: Locations outside neurons’ RF**. (b)** Example optimal solution for (L, N*) = (40, 27). Same convention as in (a). **(c)** Minimum value of cost function achieved by the optimization model with L=5 locations, plotted as a function of number of Imc-like neurons in the model; optimization was run 1000 times for each N. The minimum value progressively decreased as N increased, achieving the lowest possible value that the cost function can achieve (-L(L-1); −20 for L=5) only when N=4. In other words, the smallest number of neurons at which location-invariant selection is achievable by the model, called N*, was 4 when L=5 locations. Therefore, neuronal savings, defined as L-N*, was 1. (**d)** Fraction of optimal model solutions that had multilobed Imc neurons for all (L, N*) pairs; black bars – k_max_=3, red bars – k_max_=10. (**e)** Fraction of optimal model solutions that satisfy the “optimized lobe-overlap” property. Same conventions as in (d). **(f)** Schematic plot illustrating the need for the optimized lobe-overlap property for multilobed Imc neurons. Shown is a two-lobed Imc neuron (*middle*). When stimuli S1 and S2 are placed such that they both lie within the RF of this Imc neuron (*left*), the resulting zone of suppression generated by this Imc neuron in the OTid spares both stimuli (*right*). Thus, selection for this location-pair cannot be achieved by just this Imc neuron. **(g)** Minimum distance between neurons across model solutions and permutations recruited for solving selection for nearby location-pairs plotted as fraction of the maximum possible distance between neurons (Methods). Green dashed line: Distance cut-off for ‘distant’ neurons. **(h)** Maximum distance between neurons across model solutions and permutations recruited for solving selection for distant location-pairs plotted as fraction of the maximum possible distance between neurons (Methods). Magenta dashed line: Distance cut-off for ‘nearby’ neurons. **(i)** Fraction of optimal model solutions that satisfying the extensive intersection property. Same conventions as in (d). **(j)** Histogram of distance between centers of RF lobes within individual multilobed neurons for a randomly selected model solution for (L, N*,k_max_) = (40, 27, 3). *Inset*: Maximum elevational distance between lobes of a multilobe neuron for the same model solution. Lobes of neurons in model solutions were arbitrarily placed and widely spread. * indicates mean significantly different from 0 (p<0.001); one tailed t-test. **(k)** All 24 possible permutations of the model solution in Fig. 4a; same conventions as in Fig. 4a). For each permutation, there is at least one pair of nearby neurons that encode distant locations (indicated by blue box).

**Fig. 6.**
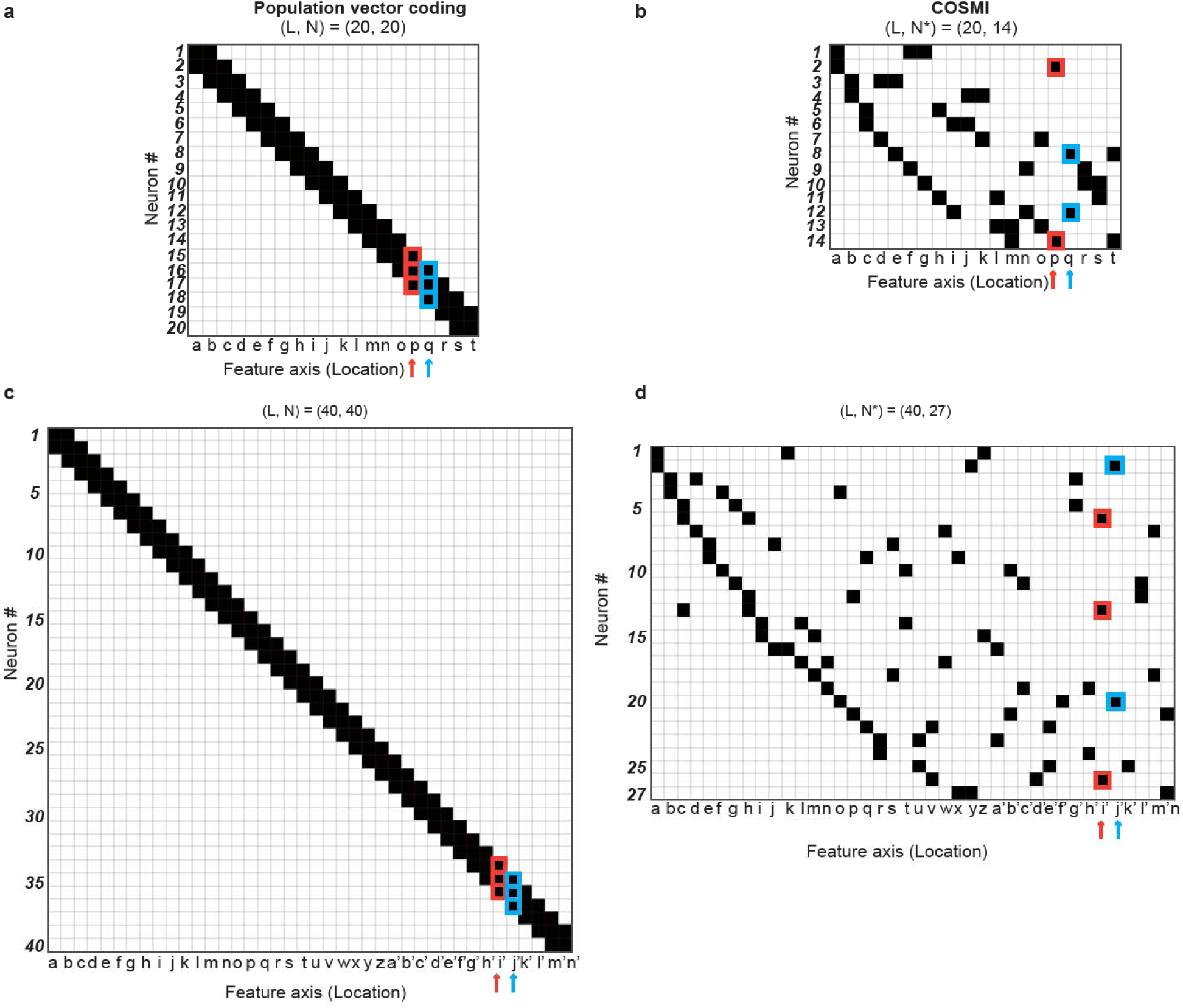
Conceptual differences between traditional population vector coding of space versus COSMI coding of space. Related to Fig. 4. **(a)** Population vector coding. Schematic illustration (heat map) of the RFs of 20 single-lobed neurons with overlapping RFs, encoding 20 feature values (say, locations). Neurons are numbered from 1 to 20 (rows), locations are denoted by alphabets (a to t; columns). Black indicates the locations at which each neuron is active. The RF of a given neuron (row) can be read out by looking at the black pixels along that row; the neurons activated by a stimulus at a particular location (column) can be read out by looking at the black pixels along that column. It is evident that each stimulus at a particular location Fig. 1. **Visual receptive fields (RFs) of Imc neurons have multiple distinct firing fields (‘lobes’). (a)** 2-D visual RF of Imc neuron: raster plot of neuron’s responses to visual stimulus presented at different spatial locations. *<u>Inset-top</u>*: Gray line – stimulus onset; red lines – time window used to calculate firing rate; evoked firing rates in Imc were high (median = 76.5 Hz; n=114 neurons). *<u>Inset-bottom:</u>* Average spike waveform for neuron in **a**; identified as high-quality unit (Methods); mean (black) ± S.D (gray). **(b)** Color coded firing rate map corresponding to **a**. **(c)** Rate map in **b** re-plotted as distribution of points in a 2-D plane and subjected to spatial clustering (Methods). Shown are the best single (*top-left*), best two (*top-right*), and best three clusters (*bottom-left*) fitted to the data using the density peaks clustering method^19^ (Supplementary Fig. 2c; Methods). *Bottom-right*: Plot of GAP statistic, a robust model selection metric, against the number (*k*) of clusters fitted to data^20^ (Methods). Red point: statistically optimal and signature property #3 – Figs. 4 and 5). (Note: in A, the maximum number of pixels in a neuron’s RF was chosen to be 3, to match the maximum number of lobes in the RFs of multilobe neurons in B.) **(c)** Another illustration of population vector coding using overlapping single-lobed RFs, but with 40 locations and 40 neurons; conventions as in A. **(d)** Another illustration of COSMI with an optimal model solution using overlapping multi-lobed RFs, but with 40 locations and 27 neurons (N*=27 neurons for L=40 locations; see Fig. 4 and text). Conventions as in (b).

